# Mapping cellular-scale internal stiffness in 3D tissues with smart material hydrogel probes

**DOI:** 10.1101/840736

**Authors:** Stephanie Mok, Sara Al Habyan, Charles Ledoux, Wontae Lee, Katherine MacDonald, Luke McCaffrey, Christopher Moraes

## Abstract

Local stiffness plays a critical role in cell function, but measuring rigidity at cellular length scales in living 3D tissues presents considerable challenges. Here we present thermoresponsive, smart material microgels that can be dispersed or injected into tissues and optically assayed to measure internal tissue stiffness over several weeks. We first develop the material design principles to measure tissue stiffness across physiological ranges, with spatial resolutions approaching that of individual cells. Using the microfabricated sensors, we demonstrate that mapping internal stiffness profiles of live multicellular spheroids at high resolutions reveal distinct architectural patterns, that vary with subtle differences in spheroid aggregation method. Finally, we determine that small sites of unexpectedly high stiffness (> 250 kPa) develop in invasive breast cancer spheroids, and in *in vivo* mouse model tumors as the cancer progresses towards metastatic disease. These highly focal sites of increased intratumoral stiffness likely form via active cell mechanical behavior, and suggest new possibilities for how early mechanical cues that drive cancer cells towards invasion might arise within the evolving tumor microenvironment.

## Introduction

Exquisitely structured tissues and organs arise from a homogenous blastomere through spatial patterns of cell proliferation, migration, and differentiation, in concert with matrix secretion and remodelling ^1–4^. Mechanical features of the local microenvironment are critical regulators of these cellular processes ^5–12^, and tissue stiffness is now well-established to drive fate-function relationships during development ^13,14^; disease progression ^15–18^ and tissue homeostasis ^19–21^. However, our technical ability to monitor stiffness at the cellular length scale during tissue development remains severely limited, and could be critically important in elucidating biophysical mechanisms of tissue morphogenesis and disease.

Conventional mechanical characterization techniques provide only a limited view of tissue stiffness, particularly at the “meso”-length scale of individual cells. Macroscale measurement tools such as tensional or shear rheometry cannot capture local stiffness variations around cells^22^, while high-resolution tools such as atomic force microscopy are ideally suited for sub-cellular nanoscale measurements, and are limited to measuring near-surface stiffness in two-dimensional or cut tissue sections. Although non-contact techniques such as ultrasound elastography or magnetic cytometry^23–25^ provide limited remote access to address these issues of scale, they cannot mimic a cell’s ability to integratively interrogate the surrounding tissue by applied deformations that span 10s of microns ^26,27^.

Serwane *et al.* recently developed an intriguing strategy to measure tissue mechanics with injectable, cell-sized, magnetic oil droplets, that deform in response to applied magnetic fields to quantify local tissue mechanics in soft tissues such as zebrafish embryos ^28^. This powerful approach provides unique insight into local stiffness evolution during development, but the small droplet volumes allow only very low actuation forces, and can hence only measure stiffnesses of < 1 kPa. More broadly, oil droplets also split apart during large scale morphogenesis, limiting the monitoring period; and this technique requires specialized equipment and expertise for simultaneous magnetic and optical probing, which limits experiments to small, thin, transparent, tissues that can stimulated with a uniform magnetic field.

Here, building upon recent materials-based strategies to generate local deformations within porous materials^22^, we introduce microscale temperature-actuated mechanosensors (μTAMs) that can be designed to measure a wide range of physiological stiffnesses, and embedded in engineered tissues or animal models to monitor extensive and long-term stiffness evolution at the length-scales of individual cells. μTAMs are spherical, thermoresponsive hydrogels that remain compact at tissue culture temperatures, but swell when cooled by a few degrees. Rigidity of the surrounding tissue either permits or restricts swelling (Fig. 1A), which can hence be calculated by observing the sensor expansion ratio. In this work, we first develop the design principles with which to optimize hydrogel formulations for various measurement sensitivities; and then demonstrate that μTAMs can be integrated into engineered tissues and animal models to track evolution of stiffness patterns. We demonstrate that distinct spatial patterns of stiffness arise in multicellular aggregates based on aggregation method; and that highly localized “hot spots” of considerably elevated intratumoral stiffness emerge during progression of breast cancer towards metastatic invasion.

**Figure 1.**
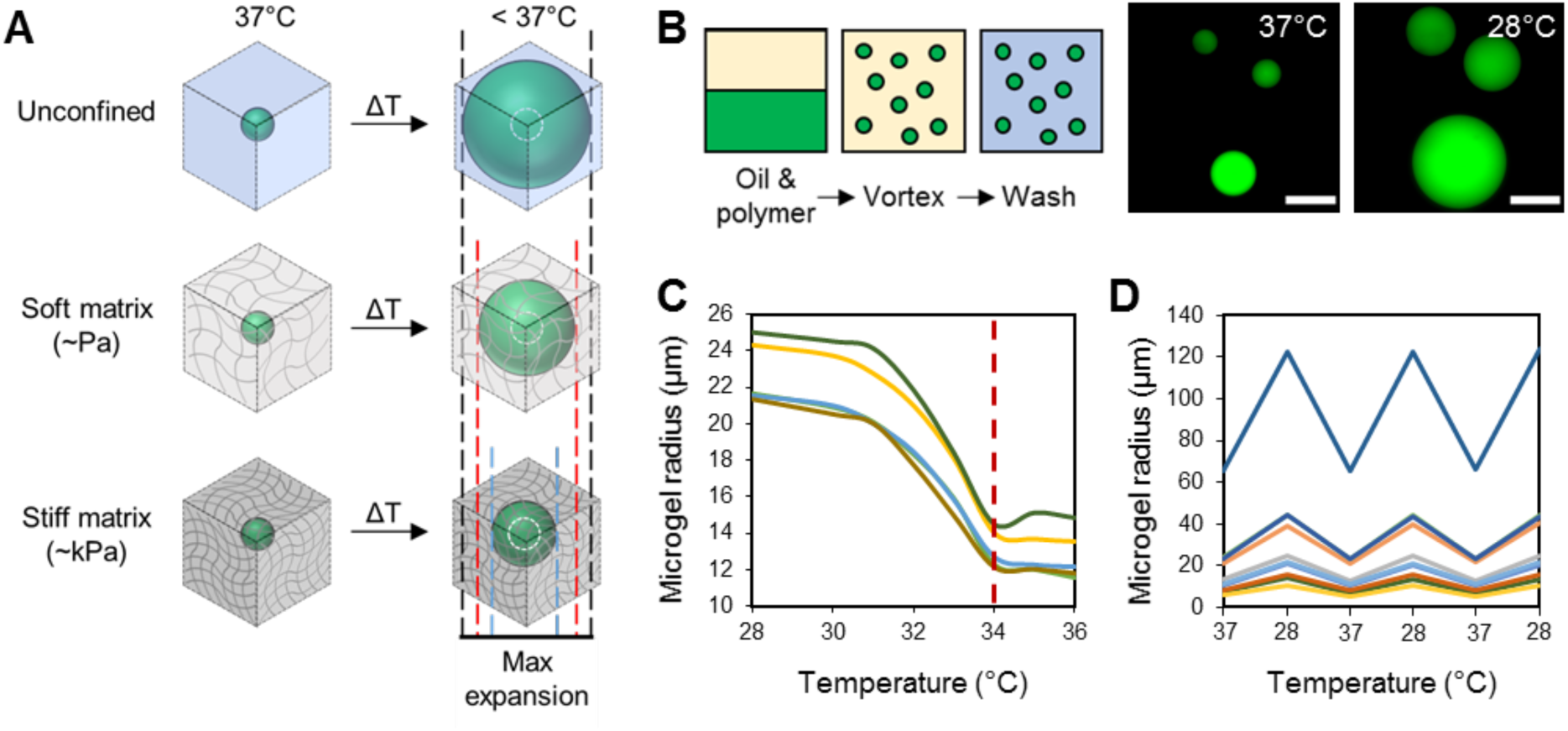
Conceptual overview of the use of thermoresponsive size-changing microgels as sensors of internal mechanical stiffness in 3D tissues. (A) Poly N-isopropylacrylamide (PNiPAAM) hydrogel droplets reversibly expand and collapse based on temperature. PNiPAAM microgels can be compacted at tissue culture temperatures of 37 °C and embedded in tissues of interest, where they will keep their contracted state while the tissue is maintained in culture conditions. Reducing the temperature below the lowest critical solution temperature triggers the microgels to expand. The degree of expansion permitted depends on the stiffness of the surrounding porous material. The expansion ratio of the sensor can hence be used to determine highly localized measurements of internal tissue stiffness, at or near tissue culture conditions. (B) To fabricate the hydrogels, an oil/water vortex emulsion technique is used to produce polydisperse spherical microscale temperature-actuated mechanosensors (μTAMS). (C) Swelling transitions between expanded and compacted states occur at 34°C, which can be (D) reproducibly observed over multiple temperature cycles. Scale bar = 50 μm.

## Results

### Design and characterization of μTAMs

Poly N-isopropylacrylamide (PNiPAAM) hydrogels are tunable, biocompatible, thermoresponsive materials that remain compact at 37 °C, but reversibly swell at slightly lower temperatures when solute interactions favour hydrophilic domains of the polymer^29,30^. To form PNiPAAM gels into μTAM probes, microspherical droplets of hydrogel formulations were polymerized with a fluorescent label^31^, in an oxygen-free, oil/water vortex emulsion (Fig. 1B). This produces polydisperse hydrogel particles with expanded diameters that range from 10 to 100 μm (Supplementary Fig. S1), which is comparable to the size and mechanical sensing range of many adherent cells ^32^. The fabricated μTAMs retain their ability to reversibly shrink above a lower critical solution temperature of ∼34 °C ^33^ (Fig. 1C, D). The thermoresponsive diameter change was independent of μTAM size, and tunable based on the hydrogel formulation (Supplementary Fig. S2). Free expansion was tunable between 1.92 ± 0.05 and 2.7 ± 0.09 for the polymer formulations tested. To confirm suitability in tissue culture conditions, we tested μTAM expansion in physiologic protein-rich conditions, as long-chain molecules in the cellular milieu may molecularly crowd and interfere with the polymer-water interactions necessary for expansion. Free expansion ratios of non-functionalized μTAMs were not significantly altered in even 100% fetal bovine serum (FBS; Supplementary Fig. S2), which contains supraphysiologically high levels of soluble protein^34^.

μTAMs require an adhesive matrix protein coating to support integration into tissues, which may impact their expansion characteristics through transport limitations or mechanically restrictions. Collagen I was selected as a candidate coating for all described experiments, as it is the most abundant matrix in the tissues studied. Standard sulfo-SANPAH crosslinking chemistry^31^ produced a monomeric collagen coating on the μTAM surface, and did not significantly affect the free expansion ratio in standard culture conditions (Supplementary. Fig. S2). We did observe a small and non-significant increase in expansion variability in the 100% FBS condition, likely arising from collagen interactions with supraphysiologically high concentrations of albumin present in FBS ^34^. Since *in vivo* interstitial albumin levels are an order of magnitude lower than in this extreme case^35^, this mechanism is unlikely to impact swelling behaviour in tissues. Together, these results confirm the suitability of PNiPAAM for repeated expansion cycles *in situ.*

### Design and characterization principles for μTAM hydrogel formulations

To select the appropriate hydrogel formulations and model deformation, we required a conceptual framework with which to design μTAMs for tissues of different rigidity ranges. Theoretically, a complete molecular simulation from first principles could determine the stored energy density of various hydrogel formulations, but such approaches would require a combination of multiscale structural, thermodynamic and molecular-interaction simulations with supporting characteristic measurements. As a first approximation, we instead reasoned that the dimensional expansion of compacted μTAMs is a balance between mechanical energy stored in the compressed sensors, and the mechanical work required to deform the surrounding material during expansion. Compacted μTAMs can hence be conceptualized as springs that are pre-loaded by thermodynamic expulsion of water prior to embedding in the tissue. Reducing the temperature releases this pre-strain, and the springs return to a new equilibrium position that is influenced by the rigidity of the surrounding material (Fig. 2A).

**Figure 2.**
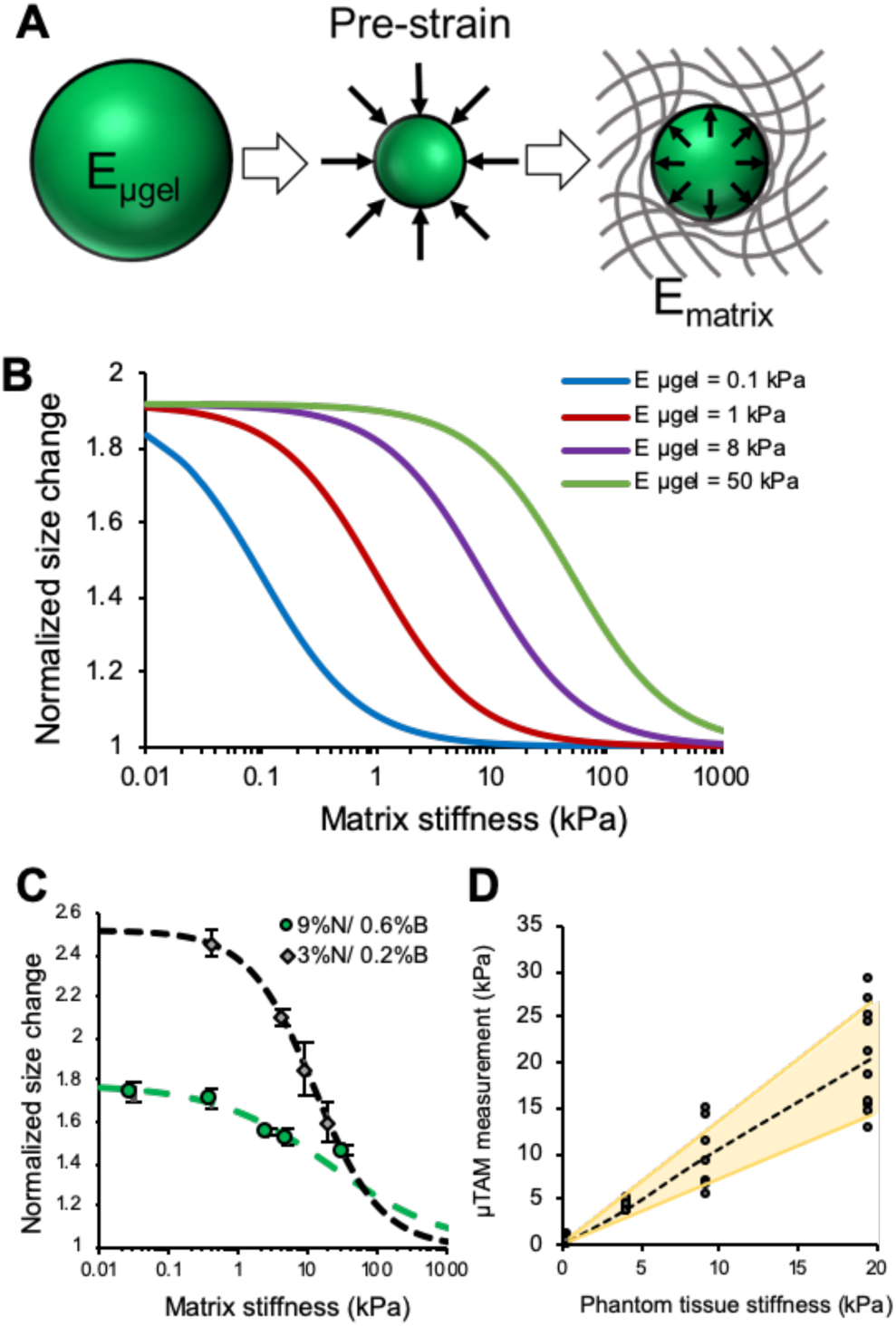
Modelling and characterization of μTAM expansion. (A) μTAMs can be modelled as pre-strained springs when compacted, which then deform the surrounding matrix when the pre-strain constraint is removed. (B) Simulations using this conceptual approach indicate that μTAMs sensitivity to the stiffness of the surrounding matrix can be tuned based on stored strain energy in the μTAM, which depends on μTAM rigidity over the actuation stroke length and applied pre-strain. A characteristic sigmoidal curve is observed with maximized measurement sensitivities in distinct measurement regimes. (C) Empirical characterization data demonstrates similar sigmoidal behaviors base on μTAM polymer formulation. Data represented as mean +/- SD (n = 6 to 11 μTAMs). Dashed line shows simulated data from a sigmoidal data fit with iteratively optimized parameters (Supplementary Table S3). (D) Multiple μTAM measurements of apparent matrix stiffness are compared against rheological measurements of matrix stiffness to determine the precision of each measurement. A linear relationship between matrix stiffness and expected measurement precision was observed and characterized.

To develop finite element computational models, we approximated the stored energy density as proportional to microgel rigidity and the degree of initial compressive pre-strain. This approximation does treat any non-linear stiffening effects as a single lumped parameter, but should still provide insight into design criteria for desirable PNiPAAM properties. Simulated spherical μTAMs of defined stiffness were isotropically pre-strained and placed within an encapsulating linear elastic material. When the pre-strain is released, a characteristic negative sigmoidal curve for μTAM expansion is produced as a function of encapsulating tissue stiffness (Fig. 2B). This is reasonable, as μTAM expansion should asymptotically approach the free expansion ratio in sufficiently soft tissues, and the completely compressed size in excessively stiff tissues. Increasing the mechanical rigidity of the μTAMs while maintaining the pre-strain levels increases the stored strain energy, shifting the sigmoidal measurement curves to provide greater sensitivities for stiffer tissues. Similar results were observed when increasing the pre-strain while maintaining μTAM mechanical rigidity. Hence, tuning the μTAM expansion ratio and mechanical rigidity can together be used to optimize stored mechanical energy in the sensors, to make measurements with desired sensitivities to tissue stiffness.

### Sensor calibration and validation in engineered tissues

To experimentally test the trends expected through simulation, we encapsulated μTAMs in tissue phantoms formed from stiffness-tunable polyacrylamide hydrogels (Supplementary Table S2). Polyacrylamide exhibits linear elastic mechanical properties over a large strain range ^31^, making it an ideal phantom material for these tests. Although the PNiPAAM formulations had similarly high rigidities in their compacted states, we were unable to independently tune the expanded stiffness and expansion ratio of the μTAMs (Fig. S3), making it difficult to predictively tune the lumped strain energy parameter underlying the model. However, low- and high-polymer content formulations were tested, and μTAM expansion in tissue phantoms followed the expected negative sigmoidal curve for increasing tissue stiffness (Fig. 2C). The sensitivity and measurement ranges were also distinct, demonstrating that μTAM formulation can be optimized for tissues with distinct stiffness regimes. We then determined the error associated with each individual μTAM measurement by comparing the μTAM-reported apparent tissue stiffness with the known stiffness of the tissue phantom, and empirically found that measurement errors can be modelled as linearly increasing with measurement values (Fig. 2D).

Finally, to verify that the μTAMs work as expected in an engineered tissue, we embedded them in multicellular spheroids (Fig. 3A, B), which are commonly used to model three-dimensional, diffusion-limited, and high-cell density tissues ^36^. Based on our calibration and modelling experiments, we elected to use the 3% NiPAAM/0.2% bisacrylamide formulation for all described experiments, as it provides greater measurement sensitivity over the stiffness range expected in spheroids ^37,38^. A model T47D cell line was mixed with μTAMs, and formed into spheroids by confinement^39^. As a first demonstration of μTAM stiffness sensing, we measured spheroid rigidities before and after tissue crosslinking through paraformaldehyde fixation, which we verified would not affect μTAM operation (Supplementary Fig. S4). Embedded μTAMs remained circular in both their compacted and expanded states, in both live (soft) and fixed (stiff) tissues, indicating that the expansion force generated is sufficiently large to overcome any small gradients of apparent stiffness that might exist around each sensor (Supplementary Fig. S4). The apparent stiffness at all sensor sites changed significantly from 0-10 kPa in live tissues to 0-46 kPa after fixation, demonstrating that the sensors function as expected (Supplementary Fig. S4). Interestingly, some sensors reported low apparent local stiffnesses even after fixation, likely because all sensors are not equally surrounded by cells within the spheroid. This heterogenous cell density is expected based on previous studies by us ^31^ and others ^40^. These results also confirm that water transport, even within densely crosslinked spheroids, is sufficient to swell the μTAMs.

**Figure 3.**
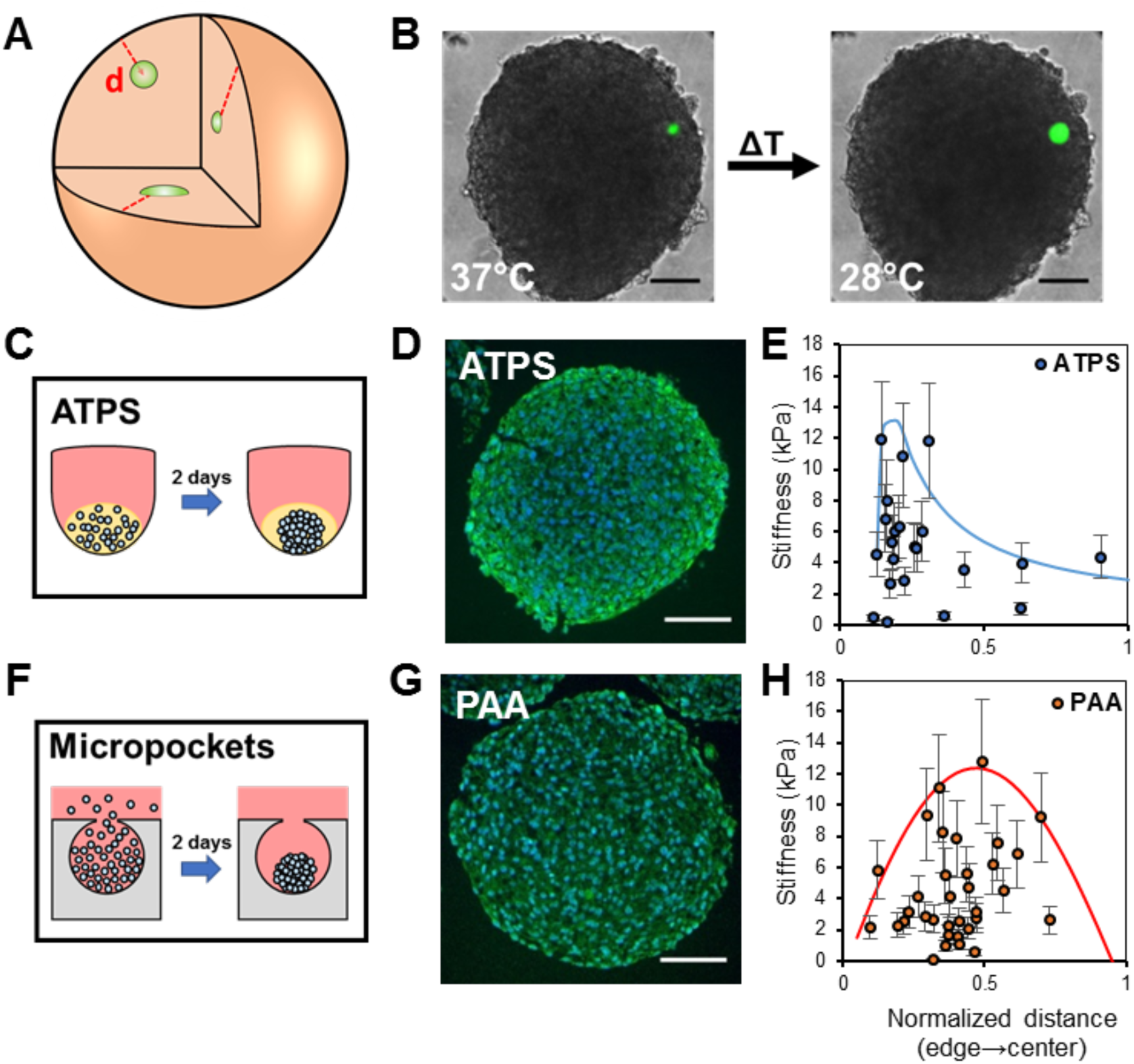
Distinct spatial patterns of internal multicellular spheroid stiffness are created based on tissue formation technique. (A) NiPAAM microgels can be randomly incorporated into 3D multicellular spheroid cultures during the tissue formation process. (B) Spheroids are formed and maintained at 37 °C, where μTAMs remain compacted. Measurements of apparent stiffness can be made by reducing the temperature and expanding the μTAMs. (C-E) Fibroblast spheroids can be formed using (C) a printable aqueous two-phase system (ATPS), in which cells are confined within a small droplet within two immiscible liquids. Alternatively (F-H), the spheroids can be formed by confining cells within a small cavity in a hydrogel where they are allowed to aggregate. (D, G) These techniques produce grossly similar spheroids, with subtle distinctions in internal architecture as assessed by tissue sectioning and staining (green = f-actin; blue = nuclei; scale bar = 100 μm). (E, H) μTAM measurements of apparent internal stiffness reveals distinct patterns of stiffness peaks through the thickness of the tissue. Data presented as measurement +/- expected error, with an enveloping inverse Gaussian distribution fit to each dataset.

### Spatial patterns of internal spheroid stiffness depends on cell aggregation method

Since spheroid architecture can be internally heterogeneous, we asked whether μTAMs can be used to map these rigidities, and whether handling methods may affect these mechanical patterns. We hence formed 400-500 μm diameter spheroids containing 1-3 μTAMs from HS-5 fibroblasts, using an aqueous two-phase system (ATPS) that confines cells to a small volume (Fig. 3C) ^41^; and a micropocket-based system in which cells passively settle into and aggregate in hydrogel cavities (Fig. 3F; Supplementary Fig S5) ^39^. These two techniques both rely on cell-driven aggregation and compaction in confined volumes, and should hence produce reasonably similar structures. No significant differences in internal cell density patterns were found in H&E-stained histology sections of the two spheroid types (Supplementary Fig. S5). However, analyzing spatially-mapped μTAM measurements revealed method-dependent patterns of apparent stiffness. Significant mechanical heterogeneity is observed across the spheroids in both cases, with measurements ranging between 0 and 12 kPa. Using an enveloping inverse Gaussian function fit to the peak values, we demonstrate that global peaks of apparent stiffness occur closer to the spheroid edges in the ATPS-formed spheroids (Fig. 3E, H), suggesting that distinct mechanical architectures arise based on even subtle differences in spheroid formation technique.

Measurements taken close to the spheroid boundaries likely under-report local stiffness, because of the geometric effects of a nearby free surface. Hence, ATPS-spheroids are likely stiffer at the tissue periphery, which drives the observed shift in peak stiffness towards the tissue edge. Observations that cell alignment is increased (Supplementary Fig. S5) and accompanied by a distinctive f-actin ring structure at the spheroid boundaries in ATPS-formed spheroids only (Fig 3D, G) supports this interpretation, as increased cortical tension in a non-linear biological material should drive an increase in cortical stiffness. The factors that underlie these distinctive patterns remain uncertain, but could arise from the small osmotic compressive pressures exerted by the dextran on the spheroids in an ATPS ^42,43^. Speculatively, these subtle stiffness differences could spatially influence cell behaviour within the spheroid, partially explaining why biological findings vary considerably between research labs that use spheroids formed via slightly different methods^44^. Specific to this work however, these experiments establish the utility of μTAMs in spatially characterizing mechanical rigidities that arise in 3D tissues.

### Internal stiffness levels of engineered tumors vary with cell type

We next asked whether μTAMs could resolve conflicting reports regarding the stiffness of metastatic and non-metastatic cancer tumors. Invasive cancer cells themselves are well-established to be more mechanically compliant than non-invasive cell types ^45^, and compliant tumors are associated with local recurrence and metastasis ^46,47^. However, clinical evidence suggests that metastatic likelihood increases with tissue stiffness ^48,15^, and external mechanical stiffness is known to promote cell migration and invasion *in vitro* ^49–51^. Other studies suggest that the internal stiffness of invasive tumors is more heterogeneous than quiescent tumors ^52^. However, these observations were made using either bulk mechanical characterization, or through surface mapping of cut tissue sections which is known to release mechanical stress ^47^. Here, we aimed to use the μTAMs to characterize the internal mechanical heterogeneity of live engineered tumors generated from differently aggressive breast cancer cell lines.

Using the micropocket-formation method, we produced similarly-sized spheroids with embedded μTAMs from human metastatic breast cancer spheroids (Fig. 4A, Supplementary Fig. S6) that we have previously established to be non-invasive (T47D) and invasive (MDA-MB-231) in collagen hydrogels over 2 days in culture^39^. The apparent internal stiffness of non-invasive spheroids varied between 0 and 12 kPa. However, the range of stiffness observed in invasive spheroids was significantly increased to 300 kPa (Fig. 4B, D). To verify that these surprisingly large readings were not the result of an outlier, histogram analysis demonstrates that a third of the measurements fall into a high-stiffness regime (Fig. 4C). These results indicate that some fraction of cells within the invasive spheroids experience high local stiffness, perhaps resolving the apparently contradictory needs for high-stiffness to prime the mechanical invasive machinery of invasive cell types, while allowing the cells to be sufficiently soft to invade through the surrounding matrix.

**Figure 4.**
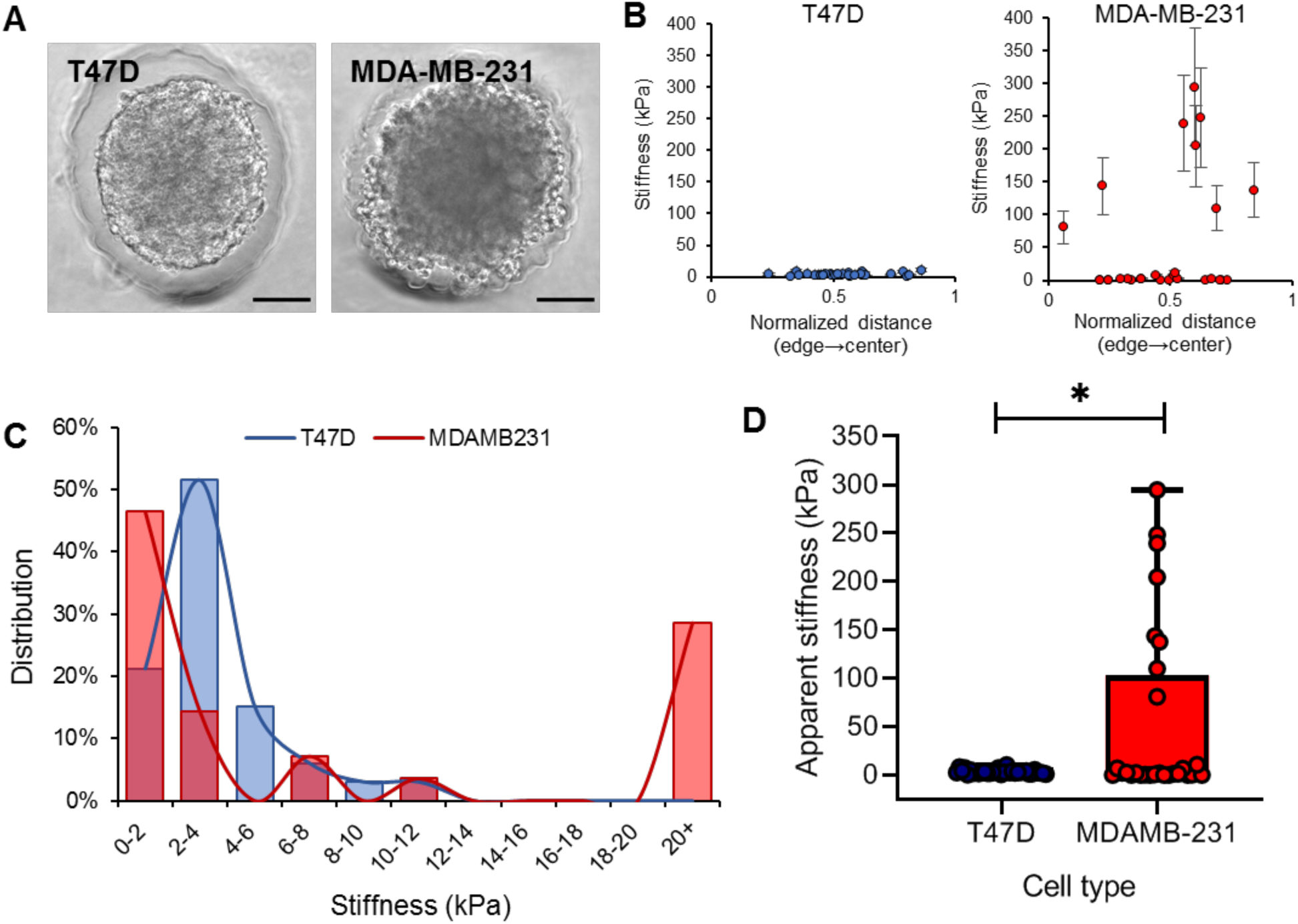
Apparent stiffness ranges within engineered tumors varies based on cell type. (A) Sample images of spheroids generated from aggressive MDA-MB-231 and less aggressive T47D metastatic breast cancer cell lines. (B) Spatial variation of stiffness in each of the tumor spheroids. Data presented as measurement +/- expected error, pooled for 1-2 μTAMs embedded in n = 25 MDAMB-231 and 27 T47D spheroids. (C) Histogram of measurement data demonstrates that a significant fraction of μTAMs in the MDA-231 spheroids registers local high apparent stiffnesses. (D) The average apparent stiffness throughout the spheroids are significantly different based on cell type. Box plots indicate the median and first to third quartile, and the whiskers span the range. * p < 0.01 (unpaired t-test, n = 33 and 28 respectively for T47D and MDA-MB-231 spheroids).

Although these findings are to our knowledge, the first *in situ* measurements of internal rigidity in live spheroids, they are consistent with several previous studies. Atomic force microscopy has been applied to measure the surface stiffness of freshly-excised metastatic tumors, which were between 0 and 16 kPa. Excision is well-established to alter the internal mechanical stress state of tissues ^47^, and punching tissue samples likely releases internal stress and disrupts the active contractility of cells that would increase stiffness in non-linear biological materials. Measurement of mechanical stiffness *in situ* has also been performed at relatively low spatial-resolution with ultrasound elastography ^53^ and mechanical stress-release analysis ^47,54^. These approaches suggest that internal tumor stiffness reach up to 150 kPa. Interestingly, these stiffnesses develop over several months, while these spheroids were formed over only two days. At these timescales, *in vivo*-like deposition of organized and stabilizing extracellular matrix would not be expected ^31^. These observations hence reflect the apparent rigidity of living cells that surround each sensor. Hence, rigidity stimuli within invasive tumors may be provided by the mechanical properties and behaviours of small groups of cells within the spheroid population.

### Long-term measurements of apparent internal tumor stiffness evolution in animal models

To extend our findings beyond *in vitro* culture, and to simultaneously demonstrate the utility of the μTAMs system *in vivo*, we injected immune-competent BALB/c mice with collagen-functionalized μTAMs and a 4T1 metastatic cancer cell line previously established to form local tumors in the mammary fat pad and spontaneously transition to invasive metastatic disease over time ^55^. 4T1 tumours have previously been mechanically characterized to be within the measurement range of μTAMs ^54,56^, and consistent with similarly-sized human tumors ^57^. Tumors were allowed to develop over several weeks. Mice were sacrificed weekly, and the excised fat pads were immediately placed in a PBS bath to control tissue temperature for stiffness measurements (Fig. 5A, B).

**Figure 5.**
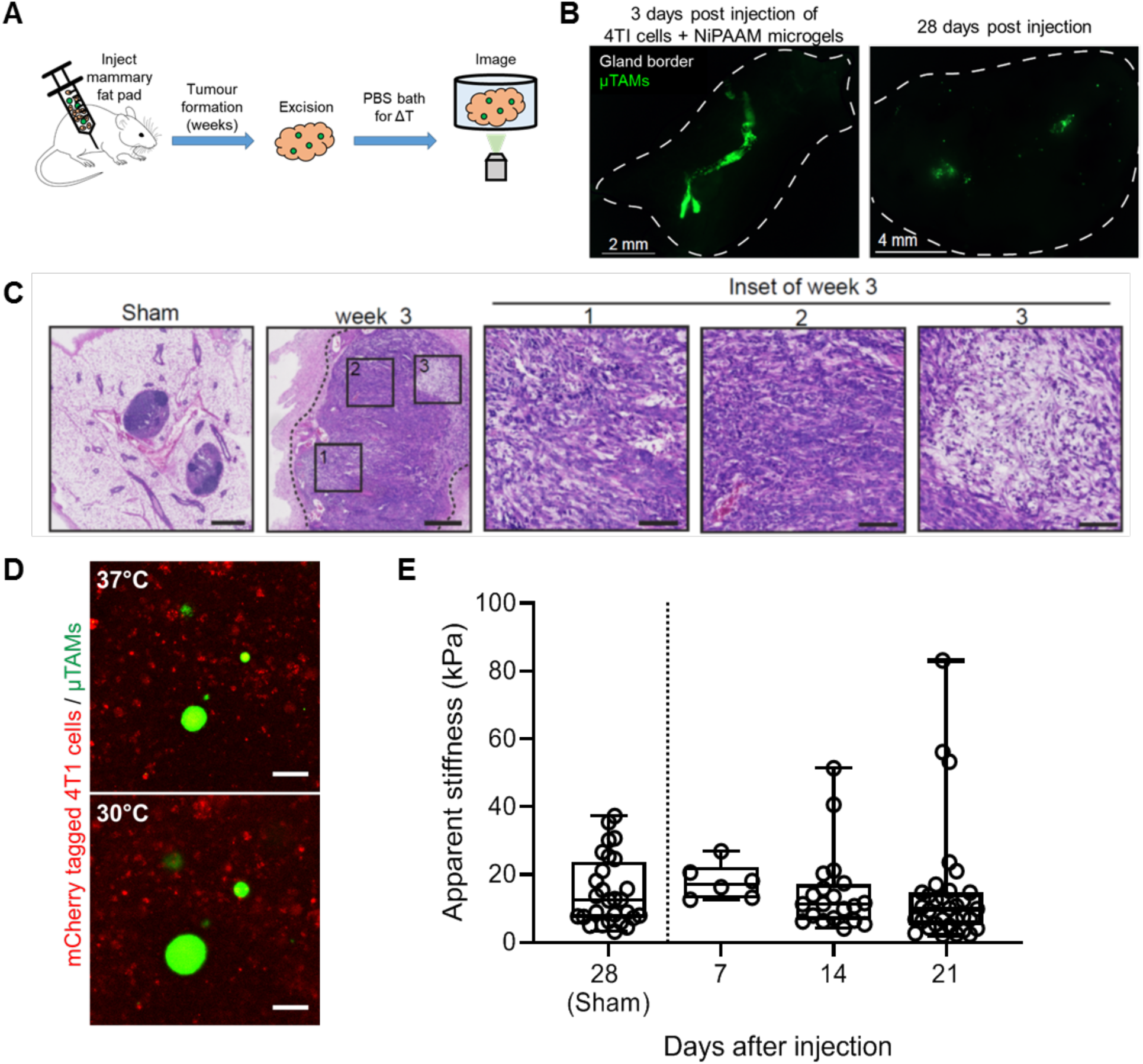
Measurements of apparent internal tumor stiffness in a metastatic breast cancer mouse model. (A) μTAMs were co-injected with mCherry-labelled T41 breast cancer cells into the mammary fat pads of female mice, and allowed to form tumors over several weeks. At various time points, tumors were excised and imaged in a temperature-controlled saline bath. (B) μTAMs are initially clustered together after injection, but disperse as the tumor develops over several weeks. (C) H&E stained tissue sections of excised fat pads at week 3 shows recovery of normal tissue architecture immediately around the needle injection site (sham, no cancer cells), and an absence of normal architecture in the tumor model. (Insets) Considerable variability in tissue cellularity is observed in distinct regions of the tumor after 3 weeks. (D) μTAMs are interspersed with mCherry-labeled 4T1 cells in the mammary fat pad, and change size when the temperature is decreased (Scale bar = 50 μm). (E) Comparison of apparent stiffness within tumors indicates an increasing number of high-stiffness outliers as the cancer progresses towards metastasis. Box plots indicate the median and first to third quartile, and the whiskers span the range.

Tumours grew rapidly in the mammary fat pad, degrading mammary gland tissue architecture by replacing fat tissue and lymph nodes with solid tumour. By week 3, a heterogeneous architecture indicative of local invasion was observed (Fig. 5C; Supplementary Figure S7). μTAMs at these later stages were diffusely distributed in the tissue, compared to tightly clustered regions seen along a well-defined wound track only 3 days after injection (Fig. 5B). No signs of additional fibrosis or inflammation were observed between sham animals injected with PBS only, and those injected with PBS and μTAMs, suggesting excellent biocompatibility of the μTAMs (Supplementary Figure S8). Only those μTAMs away from the excision wound edge were selected for analysis, to avoid measurements in regions affected by tissue stress release. These sensors were fully incorporated into the tumour tissue and retained their ability to swell and compact with temperature changes (Figure 5D).

Although the median measurements of internal tumor stiffness did not change significantly over 21 days, there was significantly greater heterogeneity observed as the tumor grew and progressed towards metastasis, with some sites stiffening to over 25 kPa (Fig. 5E). Discrete regions within tumors emerge as being much stiffer than the overall tumour as early as Day 14, which matches both our findings that local mechanical heterogeneity increases in invasive engineered tumors (Fig. 4), and observations of increasing architectural heterogeneity and stromal organization as the tumor progresses (Fig. 5C). These results demonstrate that significant mechanical heterogeneity arises over tumoral growth in vivo, suggesting a link between highly localized mechanical properties within the tumor and disease progression towards metastasis. More broadly however, in diseases such as cancer where only a few aggressive cells are required to initiate metastasis, the ability afforded by μTAMs to study highly localized mechanical microenvironments could ultimately provide an improved understanding of subpopulation-driven transitions between quiescent and malignant tumors.

## Discussion

μTAMs provide a unique opportunity for high spatial-resolution measurements of long-term stiffness evolution in a wide range of tissues. Furthermore, the tunable size and sensitivity of this general strategy allow interrogation of mechanical rigidity at multiple length scales relevant to that of a biological cell, allowing scientists to obtain a better understanding of the local microenvironment influencing a cell within a real tissue. The proof-of-concept experiments developed here together indicate that at the length scale of individual cells, the 3D tumor microenvironment is far more heterogeneous than generally expected, and our findings together suggest that microscale ‘hot spots’ of rigidity develop as metastatic tumors progress towards an invasive phenotype. Given the well-established sensitivity of cancer cells to rigidity of the local microenvironment ^49,58,59^, these studies indicate that fine spatial resolution is necessary to describe the mechanical evolution of tumors as diseases progress.

μTAMs also present some limitations that require careful consideration. First, this technique is unable to resolve time-dependent mechanical properties, such as viscoelasticity^60^, as reaching thermal equilibrium throughout the tissue makes it difficult to establish the time scales over which force is applied. Second, μTAMs may be sensitive to confounding local factors such as pH. This is unlikely to affect the present experiments, as PNiPAAM does not behave significantly differently between pH 5-8 ^61^, and tumors have internal pHs between 7.0 and 7.2 ^62^, but should be considered carefully for other tissues. Third, the requisite thermal cycling could itself influence tissue stiffness^63^. Although previous studies have demonstrated that cellular rigidity is not significantly affected between 21°C and 37°C ^64^, repetitive expansion of the sensors may theoretically induce local structural changes via damage mechanisms. To mitigate these issues in this study, we only make single measurements from each μTAM, and additional studies would be required to determine if local stress changes affect the biological systems. Fourth, the presence of these sensors itself may affect cell behaviour, as they do provide a foreign, hard surface in their compacted state. In our experiments, the hard surface presented by compacted μTAMs recapitulate microcalcification that occurs naturally in breast cancer^65^. Hence the differential responses between tissues and across timepoints in our experiments still allows us to conclude that focal stiffening is associated with invasive phenotypes. More broadly, the ability to functionalize the surface with candidate matrix molecules provide further opportunities to minimize any foreign body response.

We envision broad utility for this technology in understanding cell-scale stiffness evolution in tissues, particularly given some simple future design modifications. Developing polymer engineering strategies to tune stored strain energy through independent manipulation of expansion ratio and sensor stiffness would enable precise manipulation of actuation force and stroke length, to better simulate mechanical interrogation by specific cell types. While thermal activation was a relatively easy first step, other smart material triggers may be introduced that are faster and less disruptive, particularly for in vivo imaging. Finally, incorporating alternative imaging agents such as MRI or X-ray contrast agents would facilitate deep tissue imaging, allowing us to develop a tissue-scale cellular perspective of the local mechanical microenvironment during the highly complex processes of development and disease progression.

## Methods

All methods have been described in detail in Supplemental Information.

## Acknowledgements

We thank Profs. Richard Leask and Milan Maric for rheometer access and expertise; the Goodman Cancer Research Centre Histology Facility for assistance with processing, embedding and section tissue samples; and Lisa Zhao for schematic figures on spheroid culture formation. This work was supported by the Canadian Cancer Society (Grants # 704422, 706002) and the Canadian Institutes for Health Research (Grant # 01871-000) to C.M. and L.M., and the NSERC Discovery RGPIN-2015-05512 (C.M.). We gratefully acknowledge support from NSERC Canada Graduate Scholarship-Doctoral to S.M, Postgraduate Scholarships-Doctoral to W.L, and the Canada Research Chairs in Advanced Cellular Microenvironments to C.M.

## Author Contributions

S.M and C.M. formulated the idea behind the study. S.M., L.M, and C.M. designed experiments. S.M. and K.M conducted μTAM empirical characterization. S.M performed experiments in spheroid cultures, and conducted stiffness analysis for all experiments. C.L. performed finite element simulations. S.A. performed μTAM injections into mice and imaging of ex vivo tumors. S.M. and W.L. performed mechanical characterization and data analysis/fitting. L.M., and C.M. provided reagents, materials, animals, and analysis expertise. S.M and C.M. drafted the manuscript. All authors edited the manuscript.

## Data Availability Statement

Data supporting the findings of this manuscript are available from the corresponding author upon reasonable request.

