## Supplementary for "Mapping cellular-scale internal stiffness in 3D tissues with smart material hydrogel probes"

### Supplemental Material

#### Supplemental Figures

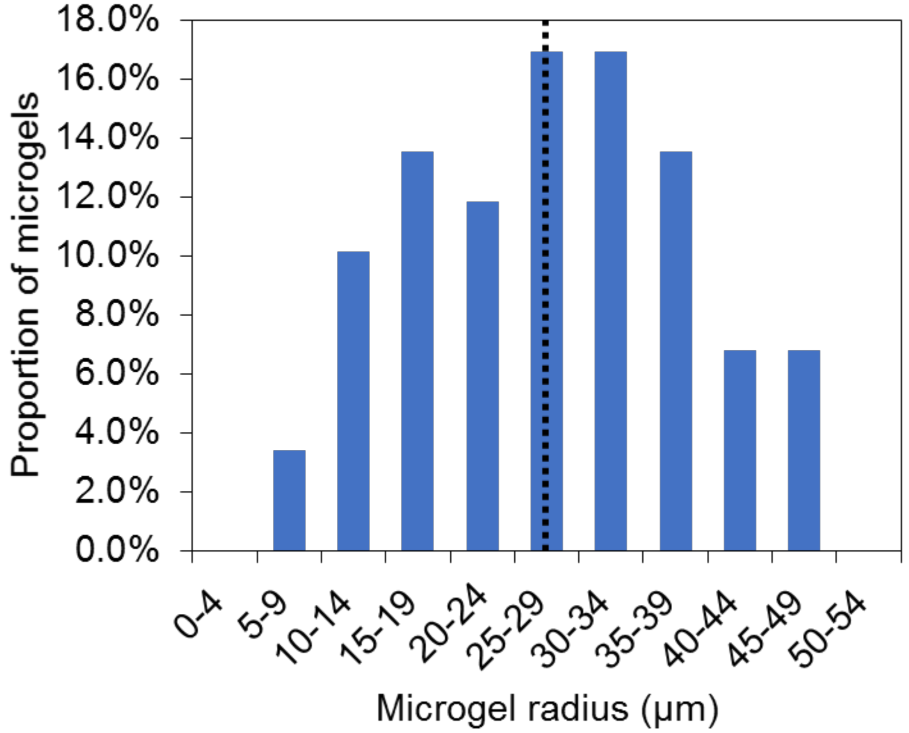


**Figure S1. NiPAAM microgel size distribution at room temperature.** Polydispersed microgels fabricated using the oil/water emulsion technique show a normal distribution of sizes with a mean radius (dotted line) of 27±11 µm. Data reported for µTAMS in expanded state (n=59).

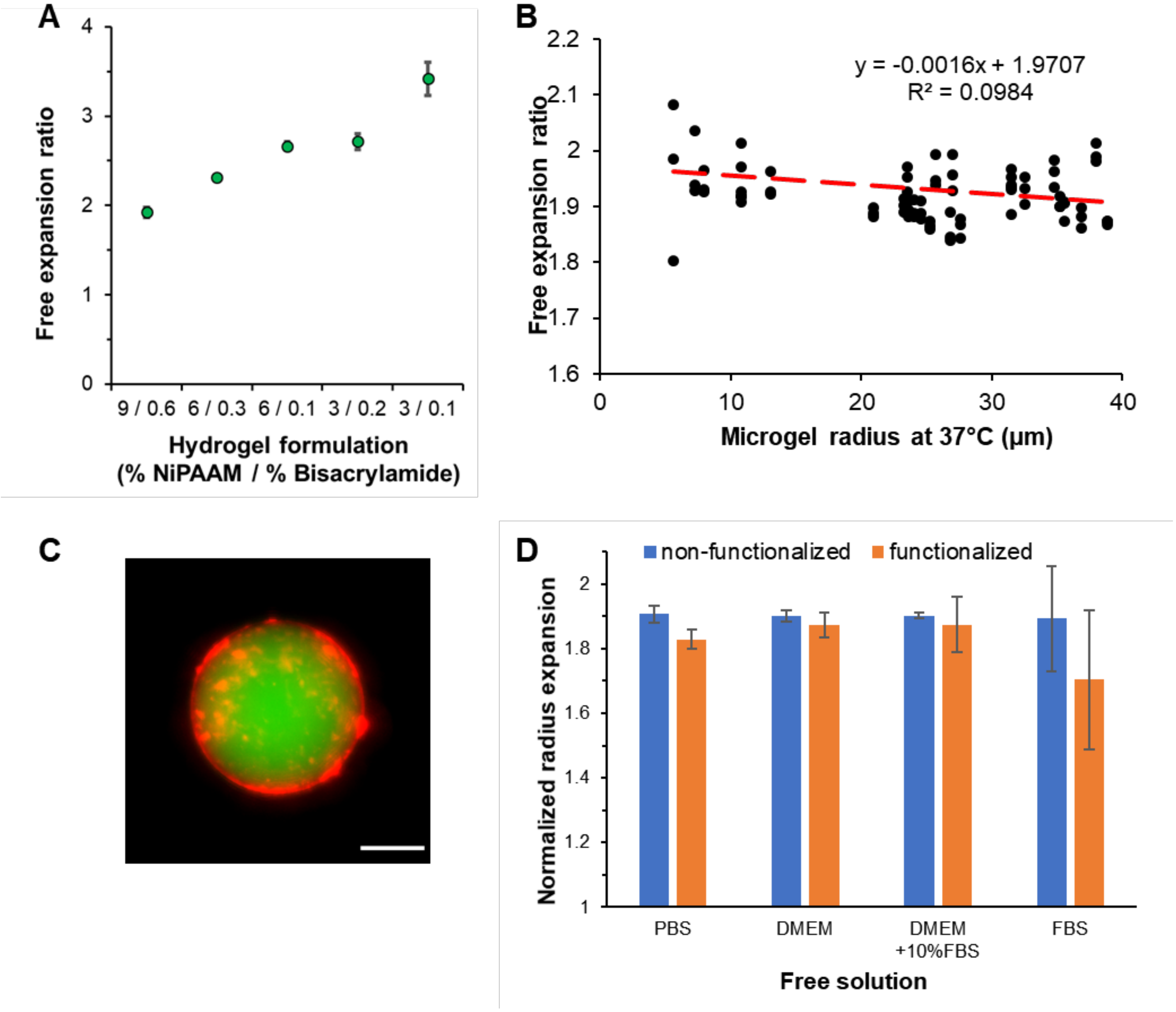


**Figure S2. Unconfined thermoresponsive free expansion of µTAMS in solution.** (A) The free expansion ratio between the measured diameters of microgels in their expanded and contracted state varies for various tested hydrogel formulations (n= 8-11). (B) Free expansion ratios are independent of microgel size (data presented for PNiPAAM formulation 9N0.6B (see Suppl. Table S1). Red dashed line indicates linear trend where y= 0.-0011x +1.9362, $R^2$ = 0.0468; n = 26). No correlation was observed between PNiPAAM microgel size and expansion ratio. Increased variability for smaller microgel sizes can be attributed to increased measurement error percentage under the selected imaging conditions . (C) Representative image of a µTAM (green) functionalized with Type I collagen (red). Scale bar = 25 µm. (D) Comparison of free expansion ratios for native and surface functionalized microgels in solutions with varying fetal bovine serum (FBS) protein content. Functionalization with collagen I has no statistically significant effect in PBS, DMEM, complete media (DMEM + 10%FBS), or 100% FBS, but some increased variability is observed in protein-rich suspensions. (Data presented as mean ± standard deviation, statistical analysis conducted by two-way ANOVA, not significant p = 0.56, 0.47 and 0.09 for the interaction between solutions and surface coating, within solutions, and within coatings; n = 3 for PBS, DMEM, DMEM+10% FBS, and functionalized gels in FBS; n= 6 for non-functionalized gels in FBS

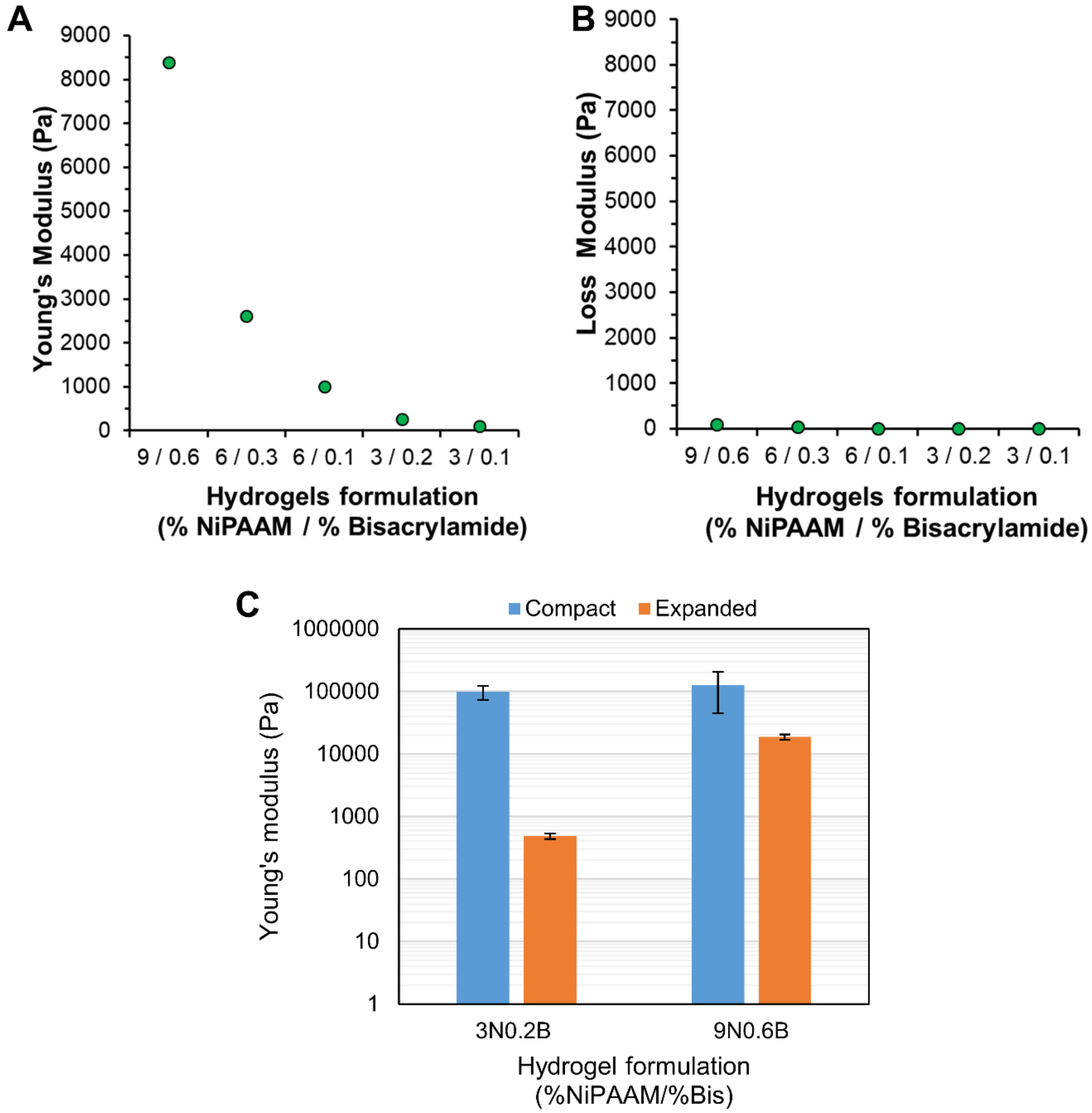


**Figure S3. Mechanical characteristics of PNiPAAM obtained via shear rheology.** (A) NiPAAM hydrogel storage modulus and (B) loss modulus. Loss moduli were minimal, indicating strongly linear elastic materials behavior. (C) PNiPAAM stiffness at the compact (>34 °C) and expanded (<34°C) hydrogel state indicate that stiffness changes dramatically between the compact and expanded state depending on PNiPAAM formualtion. Data reported as mean ± SD for n=3.

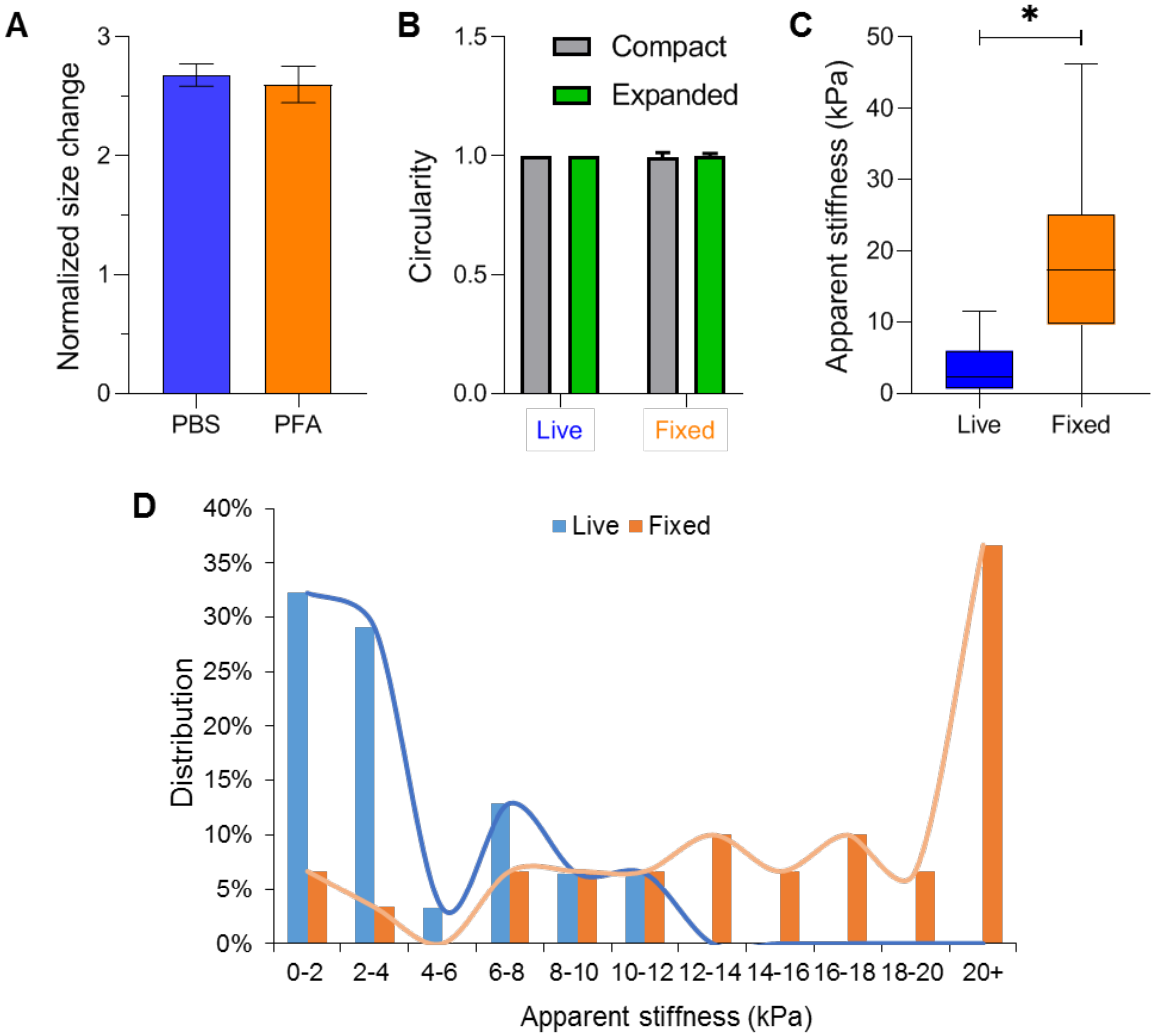


**Figure S4. Stiffness distribution within engineered T47D multicellular aggregates before and after fixation**. T-47D mammary-derived ductal carcinoma cells formed spheroids over 2 days within polyacrylamide micropockets and fixed with 4% paraformaldehyde overnight at 37°C. (A) Fixation does not affect free µTAM swelling characteristics. (n = 15 µTAMs readings for each condition). (B) Circularity of µTAMs within live and fixed tissues. No significant differences ($p = 0.061$) in circularity between compact and expanded µTAMs in either tissue condition are seen based on a two-way ANOVA. Data reported for n = 16 and 17 for live and fixed tissue condition respectively in each µTAM state. (C) Stiffness in fixed spheroids is significantly stiffer than its live pre-fixed state (n = 33 and 30 individual µTAM readings in live and fixed spheroids respectively; 30 live spheroids and 11 fixed spheroids were measured). Asterisks denotes $p < 0.0001$ according to unpaired t-test with Welch's correction. (D) Histogram of apparent stiffness distributions between live and fixed spheroids.

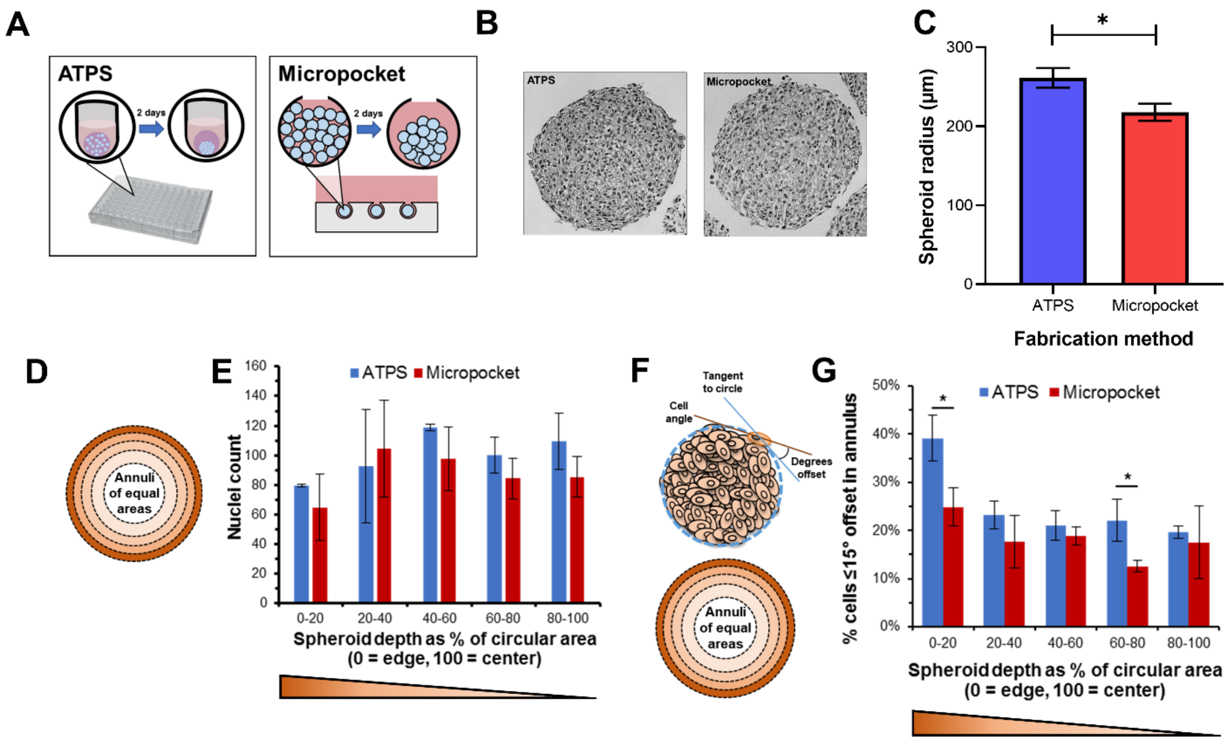


**Figure S5. Cell count and orientation in ATPS and micropocket spheroids.** (A) Schematic of spheroid formation using ATPS and micropocket techniques. (B) Representative images of H&E stained sections show that at the edge, ATPS spheroids preferentially align along the circumference of the spheroid in contrast to micropocket spheroids. (C) Characterization of spheroid dimensions formed in ATPS and micropocket systems. ATPS spheroids are slightly but significantly larger in radius than micropocket spheroids by 44 ± 4 μm (unpaired two-tailed t-test, $p < 0.0001$ for $n = 20$ and 17 respectively). (D, E) Nuclei count between the two spheroid generation methods in each annuli representing equal areas show no significant differences. ($n = 3$). (F, G) Cell orientation analysis show that cells along the periphery of ATPS spheroids preferentially elongate along the circumference of the spheroid significantly more than their micropocket counterparts. Asterisks denotes significance at $p = 0.0022$ by two-ANOVA with a Bonferonni test. ($n = 3$ spheroids for both ATPS and micropocket methods)

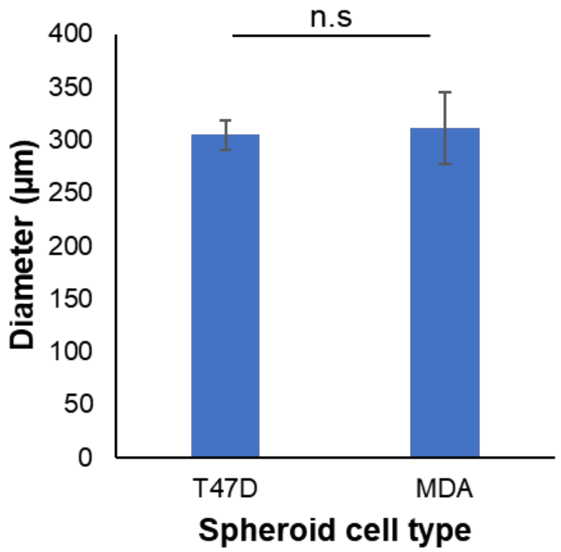


**Figure S6.** Diameter of breast cancer cell line spheroids generated using polyacrylamide micropockets. No significant difference in spheroid size between the two cell types (unpaired t-test with Welch's correction; n.s. $p = 0.383$, $n = 28$ and 26 for T47D and MDAMB-231 spheroids respectively).

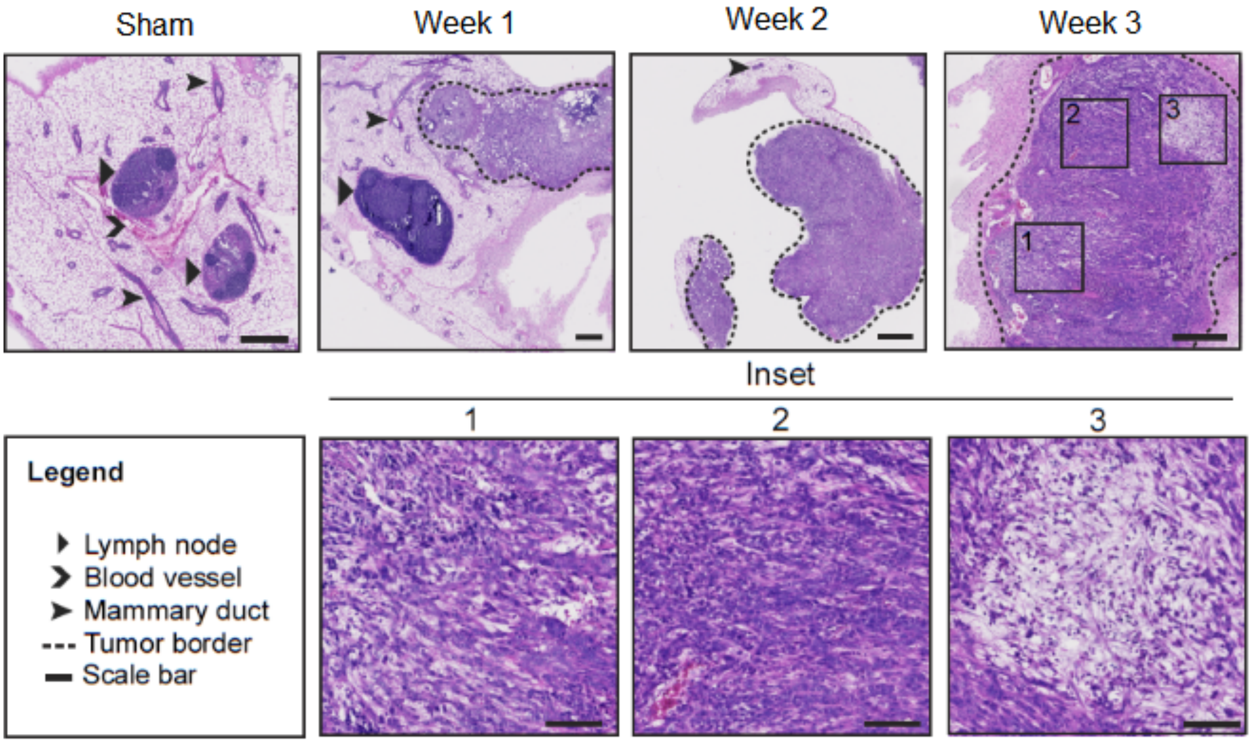


**Figure S7. Tumour development over three weeks.** Representative images of T41 tumour growth within the mammary fat pad of immunocompetent BALB/c mice. Scale bars = 500 µm. Inset scale bars 100 µm.

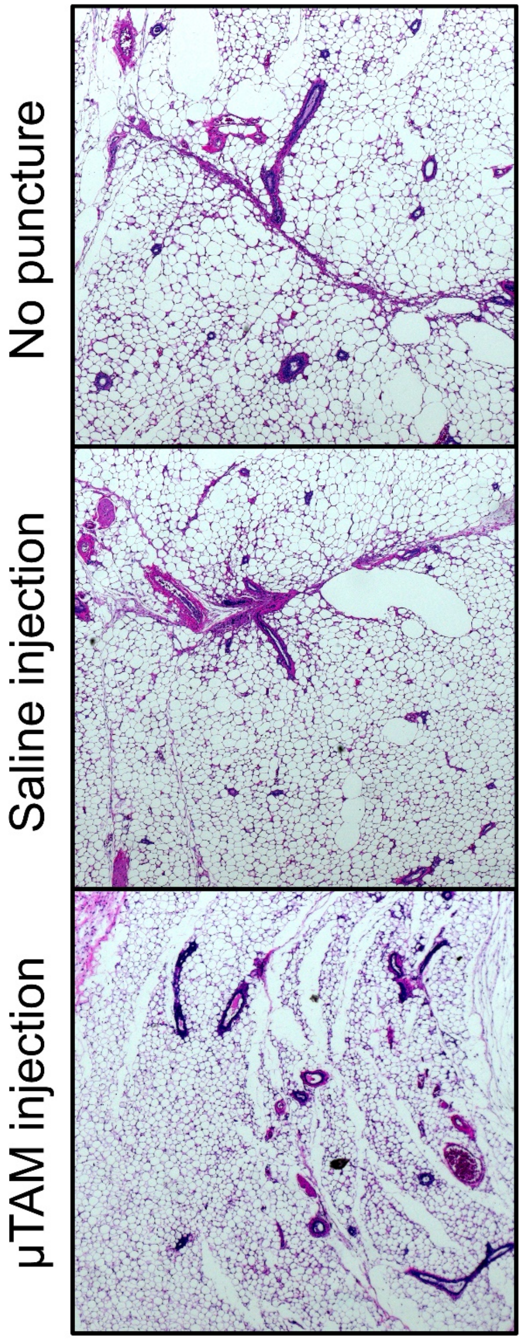


**Figure S8. Comparative histology in mouse mammary fat pads subjected to needle injection.** Representative images of mammary fat pads that were injected with saline or µTAMs using a 22G needle compared to fat pads that were not punctured show no visual differences in histology with an H&E stain. Seen in these sections are normal fat cells with occasional cross and longitudinal sections of mammary duct structures and blood vessels. Fibrotic tissue would appear as dense areas of collagen which would stain pink.

#### Supplementary Tables

**Supplementary Table S1. PNiPAAM microgel formulations.** Highlighted row in yellow indicates formulation used for all in vitro and in vivo experiments.

| NiPAAM (%) | Bis-acrylamide (%) | Formula shorthand | 20% (w/v) NiPAAM in PBS (µL) | 2% Bis-acrylmide (µL) | PBS (µL) | TEMED (µL) | Fluorescein o-methacrylate in DMSO (100 mg/mL) | 1% (w/v) APS in PBS (µL) |
|---|---|---|---|---|---|---|---|---|
| 9 | 0.6 | 9N/0.6B | 450 | 300 | 147.5 | 1.5 | 1 | 100 |
| 6 | 0.3 | 6N/0.3B | 300 | 150 | 447.5 | 1.5 | 1 | 100 |
| 6 | 0.1 | 6N/0.1B | 300 | 50 | 547.5 | 1.5 | 1 | 100 |
| 3 | 0.2 | 3N/0.2B | 150 | 100 | 647.5 | 1.5 | 1 | 100 |
| 3 | 0.1 | 3N/0.1B | 150 | 50 | 697.5 | 1.5 | 1 | 100 |

**Supplementary Table S2. Tissue phantom formulations of stiffness-tunable polyacrylamide.**

| Acrylamide (%) | Bis-acrylamide (%) | 40 % Acrylamide (µL) | 2% Bis-acrylmide (µL) | PBS (µL) | TEMED (µL) | 1% (w/v) APS in PBS (µL) | Young's Modulus based on shear rheology (Pa) |
|---|---|---|---|---|---|---|---|
| 3 | 0.05 | 75 | 24.5 | 799 | 1.5 | 100 | 150 |
| 3 | 0.11 | 75 | 53.5 | 770 | 1.5 | 100 | 400 |
| 7.5 | 0.05 | 187.5 | 27 | 694 | 1.5 | 100 | 4250 |
| 7.5 | 0.24 | 187.5 | 118 | 593 | 1.5 | 100 | 9200 |
| 12 | 0.24 | 300 | 120.5 | 478 | 1.5 | 100 | 19500 |

**Supplementary Table S3. Best fit values for PNiPAAM expansion model.**
Asterisks show empirically measured values. Bolded values are iterated best fits

| Polyacrylamide Formulation | $E_{microgel}$ (kPa) Expanded - Contracted | Free expansion ratio | B coefficient | $R^2$ fit |
|---|---|---|---|---|
| **3N0.2B** | 0.48 * - 98 * | 2.72* | N/A | N/A |
| | **12.45** | **2.52** | **0.91** | **0.94** |
| **9N0.6B** | 18.6* - 125* | 1.92* | 0.34 | 0.86 |
| | **21.60** | **1.78** | **0.514** | **0.91** |

**Supplementary Table S4. Iterated best fit values for inverse Gaussian distribution of stiffness.**

| | **Spheroid formation technique** | |
|---|---|---|
| **Parameter** | **ATPS** | **PAA** |
| **λ** | **0.115** | **1298** |
| **μ** | **2.82E+43** | **8.03** |
| **A** | **15421** | **28708** |
| **B** | **-0.113** | **7.51** |
| **C** | **716** | **-39159** |
| **$R^2$ value** | **0.85** | **0.94** |

**Supplementary Table S5. Stiffness spatial mapping data for HS-5 spheroids.**

| Fabrication method | Spheroid radius (μm) | μTAM distance from edge (μm) | μTAM radius (um) | | ratio (expanded /compact) | Apparent stiffness (Pa) |
|---|---|---|---|---|---|---|
| | | | Compact | Expanded | | |
| ATPS | 239 | 45 | 8.9 | 17.8 | 2.00 | 5968 |
| | 252 | 40 | 9.6 | 18.9 | 1.96 | 6838 |
| | 283 | 103 | 8.4 | 20.4 | 2.43 | 610 |
| | 283 | 81 | 5.8 | 11.6 | 2.00 | 6002 |
| | 265 | 208 | 6.4 | 13.3 | 2.09 | 4388 |
| | 265 | 39 | 10.4 | 18.5 | 1.77 | 11915 |
| | 256 | 160 | 9.5 | 22.5 | 2.37 | 1068 |
| | 284 | 37 | 6.9 | 14.3 | 2.08 | 4545 |
| | 284 | 50 | 7.4 | 16.4 | 2.22 | 2662 |
| | 245 | 155 | 12.1 | 25.7 | 2.12 | 3995 |
| | 256 | 53 | 15.3 | 30.5 | 1.99 | 6303 |
| | 260 | 58 | 10.5 | 23.1 | 2.20 | 2825 |
| | 255 | 42 | 9.7 | 18.5 | 1.91 | 8030 |
| | 255 | 110 | 5.6 | 12.1 | 2.15 | 3536 |
| | 264 | 31 | 6.7 | 16.4 | 2.45 | 457 |
| | 250 | 55 | 4.0 | 7.3 | 1.81 | 10882 |
| | 276 | 50 | 14.0 | 28.6 | 2.04 | 5321 |
| | 257 | 47 | 7.0 | 14.7 | 2.10 | 4270 |
| | 267 | 44 | 12.2 | 30.2 | 2.48 | 236 |
| | 268 | 69 | 7.4 | 15.1 | 2.06 | 4989 |
| | 268 | 71 | 5.4 | 11.1 | 2.06 | 4935 |
| | 264 | 82 | 14.6 | 25.9 | 1.78 | 11834 |
| Micropocket | 223 | 162 | 11.7 | 26.1 | 2.22 | 2623 |
| | 212 | 87 | 8.4 | 20.5 | 2.43 | 541 |
| | 212 | 106 | 11.8 | 24.6 | 2.08 | 4517 |
| | 212 | 134 | 18.8 | 47.5 | 2.53 | 0 |
| | 216 | 80 | 8.2 | 18.4 | 2.25 | 2241 |
| | 216 | 96 | 8.2 | 17.1 | 2.07 | 4731 |
| | 230 | 54 | 6.9 | 15.0 | 2.18 | 3127 |
| | 230 | 126 | 9.4 | 18.0 | 1.93 | 7588 |
| | 223 | 22 | 5.8 | 13.1 | 2.26 | 2141 |
| | 223 | 105 | 8.1 | 17.8 | 2.21 | 2763 |
| | 215 | 78 | 4.1 | 7.9 | 1.91 | 7854 |
| | 215 | 51 | 4.7 | 9.9 | 2.11 | 4156 |
| | 222 | 48 | 7.6 | 17.0 | 2.23 | 2525 |
| | 213 | 100 | 7.8 | 17.0 | 2.18 | 3099 |
| | 198 | 81 | 11.8 | 26.4 | 2.23 | 2547 |
| | 213 | 63 | 7.0 | 13.1 | 1.86 | 9389 |

| | | | | | |
|---|---|---|---|---|---|
| 213 | 72 | 13.1 | 23.6 | 1.80 | 11104 |
| 219 | 122 | 10.1 | 19.7 | 1.96 | 6879 |
| 219 | 72 | 10.7 | 21.8 | 2.03 | 5466 |
| 219 | 75 | 9.9 | 20.8 | 2.11 | 4160 |
| 240 | 74 | 13.9 | 32.0 | 2.30 | 1696 |
| 228 | 44 | 7.4 | 16.5 | 2.25 | 2307 |
| 228 | 112 | 7.5 | 13.1 | 1.75 | 12788 |
| 228 | 121 | 9.3 | 18.5 | 1.99 | 6221 |
| 228 | 124 | 11.7 | 34.2 | 2.92 | 0 |
| 196 | 24 | 7.1 | 14.3 | 2.01 | 5851 |
| 196 | 69 | 9.2 | 17.4 | 1.90 | 8289 |
| 212 | 77 | 6.8 | 16.3 | 2.38 | 944 |
| 220 | 71 | 12.8 | 28.4 | 2.22 | 2671 |
| 220 | 82 | 9.8 | 19.8 | 2.02 | 5561 |
| 220 | 60 | 10.1 | 25.4 | 2.51 | 49 |
| 207 | 145 | 9.7 | 18.1 | 1.86 | 9210 |
| 233 | 68 | 10.6 | 23.3 | 2.20 | 2864 |
| 233 | 95 | 13.6 | 31.6 | 2.32 | 1524 |
| 219 | 91 | 9.7 | 22.8 | 2.37 | 1105 |
| 219 | 97 | 11.8 | 26.6 | 2.27 | 2097 |
| 239 | 45 | 8.9 | 17.8 | 2.00 | 5968 |
| 252 | 40 | 9.6 | 18.9 | 1.96 | 6838 |
| 283 | 103 | 8.4 | 20.4 | 2.43 | 610 |
| 283 | 81 | 5.8 | 11.6 | 2.00 | 6002 |
| 265 | 208 | 6.4 | 13.3 | 2.09 | 4388 |
| 265 | 39 | 10.4 | 18.5 | 1.77 | 11915 |
| 256 | 160 | 9.5 | 22.5 | 2.37 | 1068 |
| 284 | 37 | 6.9 | 14.3 | 2.08 | 4545 |
| 284 | 50 | 7.4 | 16.4 | 2.22 | 2662 |
| 245 | 155 | 12.1 | 25.7 | 2.12 | 3995 |
| 256 | 53 | 15.3 | 30.5 | 1.99 | 6303 |
| 260 | 58 | 10.5 | 23.1 | 2.20 | 2825 |
| 255 | 42 | 9.7 | 18.5 | 1.91 | 8030 |
| 255 | 110 | 5.6 | 12.1 | 2.15 | 3536 |
| 264 | 31 | 6.7 | 16.4 | 2.45 | 457 |
| 250 | 55 | 4.0 | 7.3 | 1.81 | 10882 |
| 276 | 50 | 14.0 | 28.6 | 2.04 | 5321 |
| 257 | 47 | 7.0 | 14.7 | 2.10 | 4270 |
| 267 | 44 | 12.2 | 30.2 | 2.48 | 236 |
| 268 | 69 | 7.4 | 15.1 | 2.06 | 4989 |
| 268 | 71 | 5.4 | 11.1 | 2.06 | 4935 |
| 264 | 82 | 14.6 | 25.9 | 1.78 | 11834 |

**Supplementary Table S6. Stiffness spatial mapping data for breast cancer spheroids.**

| Cell Type | Spheroid radius (µm) | µTAM distance from edge (µm) | µTAM radius (um) | | ratio (expanded /compact) | Apparent stiffness (Pa) |
|---|---|---|---|---|---|---|
| | | | Compact | Expanded | | |
| MDAMB-231 | 140 | 29 | 23.4 | 57.0 | 2.44 | 527 |
| | 131 | 93 | 18.7 | 51.1 | 2.73 | 0 |
| | 128 | 31 | 22.2 | 54.9 | 2.47 | 264 |
| | 128 | 82 | 25.7 | 66.3 | 2.58 | 0 |
| | 164 | 99 | 20.6 | 22.9 | 1.11 | 204612 |
| | 144 | 49 | 22.7 | 56.3 | 2.48 | 217 |
| | 163 | 75 | 28.8 | 71.8 | 2.49 | 135 |
| | 145 | 55 | 23.5 | 53.2 | 2.26 | 2116 |
| | 142 | 70 | 27.1 | 66.0 | 2.44 | 531 |
| | 155 | 51 | 19.3 | 43.5 | 2.26 | 2163 |
| | 164 | 102 | 19.1 | 20.9 | 1.09 | 248360 |
| | 177 | 11 | 19.3 | 23.8 | 1.24 | 80910 |
| | 164 | 35 | 23.4 | 58.1 | 2.48 | 199 |
| | 183 | 94 | 18.9 | 37.3 | 1.97 | 6551 |
| | 181 | 54 | 20.1 | 47.1 | 2.34 | 1307 |
| | 167 | 116 | 23.7 | 28.1 | 1.19 | 109892 |
| | 163 | 90 | 18.4 | 20.2 | 1.10 | 239363 |
| | 178 | 40 | 20.7 | 23.8 | 1.15 | 143493 |
| | 178 | 94 | 22.9 | 51.2 | 2.24 | 2435 |
| | 140 | 34 | 19.9 | 51.7 | 2.60 | 0 |
| | 151 | 78 | 16.5 | 29.9 | 1.81 | 10664 |
| | 151 | 67 | 20.2 | 39.6 | 1.96 | 6828 |
| | 172 | 127 | 27.3 | 66.0 | 2.42 | 668 |
| | 123 | 103 | 16.8 | 19.4 | 1.15 | 137816 |
| | 166 | 112 | 21.2 | 49.9 | 2.35 | 1210 |
| | 145 | 55 | 14.3 | 31.9 | 2.23 | 2494 |
| | 159 | 111 | 23.8 | 59.4 | 2.50 | 119 |
| | 159 | 96 | 12.2 | 13.2 | 1.08 | 294621 |
| T47D | 155 | 76 | 20.1 | 44.0 | 2.19 | 3010 |
| | 155 | 122 | 27.9 | 54.0 | 1.94 | 7377 |
| | 149 | 119 | 25.0 | 59.6 | 2.38 | 943 |
| | 141 | 62 | 14.5 | 32.9 | 2.27 | 2060 |
| | 150 | 80 | 16.8 | 38.8 | 2.31 | 1634 |
| | 150 | 93 | 15.0 | 30.4 | 2.03 | 5515 |
| | 144 | 61 | 19.3 | 43.0 | 2.23 | 2527 |

| | | | | | |
|---|---|---|---|---|---|
| 144 | 50 | 10.9 | 20.7 | 1.90 | 8242 |
| 162 | 80 | 23.5 | 52.2 | 2.22 | 2608 |
| 150 | 84 | 14.3 | 29.4 | 2.06 | 4995 |
| 156 | 97 | 24.3 | 54.7 | 2.25 | 2259 |
| 156 | 96 | 14.0 | 27.1 | 1.94 | 7372 |
| 163 | 100 | 32.0 | 72.6 | 2.27 | 2062 |
| 157 | 51 | 21.7 | 52.1 | 2.40 | 802 |
| 149 | 70 | 18.6 | 40.2 | 2.16 | 3379 |
| 149 | 128 | 23.4 | 42.7 | 1.82 | 10276 |
| 156 | 88 | 17.0 | 41.7 | 2.45 | 402 |
| 157 | 70 | 17.9 | 39.6 | 2.21 | 2717 |
| 157 | 127 | 17.5 | 39.4 | 2.25 | 2255 |
| 154 | 97 | 30.7 | 71.6 | 2.33 | 1412 |
| 149 | 77 | 20.1 | 45.7 | 2.27 | 2009 |
| 166 | 65 | 15.8 | 33.5 | 2.12 | 3965 |
| 157 | 56 | 22.7 | 52.7 | 2.32 | 1515 |
| 154 | 83 | 21.7 | 45.8 | 2.11 | 4111 |
| 158 | 72 | 26.3 | 58.0 | 2.21 | 2803 |
| 160 | 73 | 22.4 | 49.6 | 2.21 | 2693 |
| 152 | 77 | 22.2 | 49.0 | 2.21 | 2780 |
| 164 | 100 | 23.7 | 49.3 | 2.08 | 4590 |
| 141 | 33 | 15.8 | 33.4 | 2.11 | 4061 |
| 155 | 87 | 25.3 | 57.8 | 2.28 | 1892 |
| 148 | 87 | 28.3 | 62.2 | 2.20 | 2897 |
| 145 | 75 | 13.0 | 28.4 | 2.18 | 3067 |
| 136 | 100 | 18.9 | 40.6 | 2.15 | 3561 |

**Supplementary Table S7. Stiffness readings from ex vivo tumours.**

| Days post-injection | µTAM radius (um) | | ratio (expanded /compact) | Apparent stiffness (Pa) |
|---|---|---|---|---|
| | Compact | Expanded | | |
| 7 | 14.7 | 25.8 | 1.75 | 12632 |
| | 13.7 | 22.8 | 1.67 | 16276 |
| | 13.2 | 19.9 | 1.51 | 26856 |
| | 9.6 | 15.6 | 1.63 | 18121 |
| | 9.3 | 14.8 | 1.59 | 20623 |
| | 7.8 | 13.6 | 1.74 | 13219 |
| 14 | 7.2 | 14.6 | 2.03 | 5390 |
| | 8.3 | 14.4 | 1.73 | 13560 |
| | 8.3 | 13.7 | 1.64 | 17500 |
| | 8.3 | 11.0 | 1.33 | 51370 |
| | 9.0 | 16.3 | 1.81 | 10693 |
| | 12.7 | 22.7 | 1.79 | 11323 |
| | 11.4 | 19.1 | 1.68 | 15835 |
| | 14.2 | 28.2 | 1.99 | 6201 |
| | 10.9 | 19.4 | 1.78 | 11620 |
| | 11.6 | 23.0 | 1.99 | 6305 |
| | 10.3 | 20.3 | 1.98 | 6427 |
| | 15.8 | 28.2 | 1.79 | 11399 |
| | 18.8 | 39.7 | 2.11 | 4102 |
| | 8.7 | 16.6 | 1.91 | 7964 |
| | 16.3 | 31.8 | 1.95 | 7096 |
| | 9.5 | 17.5 | 1.84 | 9853 |
| | 10.1 | 15.9 | 1.58 | 21180 |
| | 14.4 | 23.0 | 1.59 | 20300 |
| | 7.4 | 10.3 | 1.39 | 40652 |
| | 11.6 | 19.9 | 1.72 | 13952 |
| 21 | 9.8 | 18.0 | 1.83 | 10007 |
| | 8.9 | 16.3 | 1.84 | 9887 |
| | 9.8 | 21.9 | 2.23 | 2479 |
| | 9.1 | 20.4 | 2.24 | 2340 |
| | 13.2 | 29.1 | 2.21 | 2749 |
| | 7.5 | 16.0 | 2.15 | 3484 |
| | 9.0 | 15.8 | 1.76 | 12359 |
| | 11.8 | 20.0 | 1.70 | 14707 |
| | 12.7 | 26.7 | 2.09 | 4368 |
| | 17.5 | 34.4 | 1.97 | 6596 |
| | 14.3 | 30.1 | 2.11 | 4165 |
| | 7.8 | 14.1 | 1.81 | 10744 |

| | | | | |
|---|---|---|---|---|
| | 8.0 | 16.0 | 1.99 | 6304 |
| | 12.0 | 18.6 | 1.54 | 23651 |
| | 16.5 | 32.3 | 1.96 | 6812 |
| | 14.1 | 27.7 | 1.96 | 6815 |
| | 7.2 | 14.0 | 1.95 | 6963 |
| | 11.5 | 22.1 | 1.93 | 7531 |
| | 14.0 | 24.6 | 1.75 | 12605 |
| | 11.4 | 23.3 | 2.04 | 5350 |
| | 8.6 | 16.0 | 1.86 | 9385 |
| | 11.5 | 21.0 | 1.82 | 10458 |
| | 8.3 | 17.2 | 2.06 | 4958 |
| | 10.9 | 24.5 | 2.25 | 2245 |
| | 7.9 | 13.6 | 1.72 | 13786 |
| | 12.3 | 21.4 | 1.74 | 13271 |
| | 5.4 | 8.6 | 1.58 | 21045 |
| | 22.4 | 29.3 | 1.31 | 56154 |
| | 14.2 | 17.5 | 1.23 | 83094 |
| | 24.8 | 32.7 | 1.32 | 53254 |
| | 11.3 | 19.2 | 1.70 | 14683 |
| | 10.9 | 18.0 | 1.65 | 17081 |
| | 10.3 | 17.4 | 1.69 | 15250 |
| 28 (Sham) | 10.8 | 20.5 | 1.90 | 8273 |
| | 7.4 | 10.4 | 1.41 | 37274 |
| | 7.1 | 10.3 | 1.46 | 30746 |
| | 12.9 | 19.7 | 1.53 | 24584 |
| | 6.6 | 12.4 | 1.88 | 8841 |
| | 6.3 | 10.0 | 1.58 | 21148 |
| | 10.0 | 20.4 | 2.05 | 5164 |
| | 9.8 | 17.2 | 1.75 | 12615 |
| | 5.7 | 11.0 | 1.92 | 7715 |
| | 8.1 | 15.7 | 1.93 | 7575 |
| | 12.2 | 21.1 | 1.73 | 13667 |
| | 11.6 | 16.6 | 1.42 | 35474 |
| | 13.4 | 20.2 | 1.51 | 26629 |
| | 8.7 | 12.8 | 1.47 | 30096 |
| | 9.9 | 19.2 | 1.94 | 7209 |
| | 7.8 | 15.6 | 2.00 | 5994 |
| | 8.6 | 14.5 | 1.68 | 15825 |
| | 11.1 | 19.5 | 1.75 | 12766 |
| | 10.1 | 20.1 | 1.99 | 6296 |
| | 11.5 | 21.5 | 1.87 | 8888 |
| | 6.4 | 13.9 | 2.17 | 3209 |

| | | | | |
|---|---|---|---|---|
| | 11.4 | 20.1 | 1.76 | 12456 |
| | 13.9 | 21.2 | 1.52 | 25336 |
| | 11.8 | 22.6 | 1.91 | 7991 |
| | 5.1 | 10.6 | 2.06 | 4918 |
| | 5.1 | 10.7 | 2.09 | 4438 |
| | 13.0 | 21.8 | 1.69 | 15443 |
| | 8.5 | 13.9 | 1.63 | 18133 |

#### Online Methods

Unless otherwise stated, all cell culture materials and supplies were purchased from Fisher Scientific (Ottawa, ON), and chemicals from Sigma Aldrich (Oakville, ON).

**µTAMs fabrication**

Separate solutions of 6% (w/v) polyglycerol polyrincinoleate surfactant (PGPR 4150; Palsgaard, 90415001) in kerosene; 1% (w/v) ammonium persulfate (APS) in phosphate buffered saline; and a prepolymerized PNiPAAM solution following Table S1 (excluding 1% APS) were each prepared in individual glass test tubes with 1-2 mL of each solution in their respective tubes. Volumes within the test tubes are fairly flexible, provided there is a matched or excess volume within the kerosene tube to create a bath. A magnetic stir bar was placed within the kerosene test tube. To purge the system of oxygen, a rubber septum stopper were used to seal each tube and nitrogen gas was bubbled through each liquid for at least 20 minutes using a 25G non-coring needle, with a second needle to vent the tubes to atmosphere. Microspherical gels were formed by drawing the desired amount of 1% APS solution into a syringe and dispensing it into the sealed test tube containing PNiPAAM components. The mixture was immediately vortexed and transferred into the kerosene bath with another syringe. An emulsion was made by vortexing the kerosene/PNiPAAM mixture for 5 to 10 seconds. Droplets were prevented from coalescing by gentle magnetic stirring for 20 minutes as the µTAMs polymerized. To facilitate washing and recovery of the µTAMs, the emulsion was aliquoted into several 1.5 mL microcentrifuge tubes. Each washing step included centrifugation at 14,800x g for 3 minutes, supernatant aspiration and µTAM resuspension with the appropriate solution. The µTAMs were first washed with fresh kerosene three times to remove the PGPR4150 surfactant, and then with PBS three times to recover the microgels in an aqueous phase. Finally, µTAMs were stored at 4°C in PBS overnight to allow gels to swell to equilibrium before further use.

**µTAMs surface functionalization**

µTAMs were suspended in a 0.05 mg/mL solution of sulfoSANPAH (GBiosciences # BC38) in PBS and irradiated under 36 W UV light for 4 minutes. The solution was aspirated and the µTAMs were washed once with PBS before being incubated in 0.05 mg/mL solution of collagen I (Advanced Biomatrix PureCol #5005B) in PBS overnight at 4°C. Gels were then washed with PBS and stored at 4°C. Prior to embedding or injection into tissues, µTAMs were UV sterilized for 45 minutes (36W UV source).

**Stiffness-tunable tissue phantoms**

Polyacrylamide hydrogels were fabricated on glass coverslips with embedded µTAMs to calibrate sensor measurements. Hydrogel-releasing hydrophobic glass slides were prepared by coating RainX onto 75 x 50 mm glass slides. Glass coverslips were silanized to bind polyacrylamide by immersion in a 0.4% 3-(Trimethoxysilyl) propyl methacrylate (MPS) in acetone for 5 minutes, washed with fresh acetone for 5 minutes, and air dried.

To embed µTAMs into polyacrylamide tissue phantoms, polyacrylamide pre-gel solutions were made according to Table 2 with a small volume of PBS replaced by an equal volume of µTAMs in

PBS. The complete pre-gel solution with NiPAAM microgels was pipetted onto a hydrophobic glass slide in multiple 127 µL drops to produce a 0.5 mm thick gel when a silanized 18 mm round coverslip was place on top of each drop. The solution was left to polymerize on a slide warmer set to 45°C for 10 minutes. This ensures that µTAMs enter the tissue in their compacted state. After polymerization, the coverslips with the attached hydrogel were peeled off the glass slide with tweezers and placed in a multi-well plate. All hydrogels were washed 3 times with PBS and left to equilibrate overnight in a 37°C incubator before thermal cycling and imaging.

**Cell culture**
Human HS-5 fibroblasts (ATCC CRL-11882); and T47D (ATCC HTB-133) and MDAMB-231 (ATCC HTB-26) breast cancer cell lines were cultured in Dulbecco's modified eagle media with 10% fetal bovine serum (FBS) and 1% anti/anti (complete media). Cells used for mice experiments were Mouse 4T1 (ATCC CRL-2539), which were cultured in RPMI 1640 (Wisent) with 10% FBS, 1% sodium bicarbonate, 0.5% sodium pyruvate and 0.5% HEPES. When the cells reached at least 80% confluence (70% for 4T1 cells to maintain tumorigenic characteristics), they were detached using 0.25% trypsin-EDTA and either subcultured into a new culture vessel at a 1:10 ratio or used as a single cell suspension for experiments.

**Spheroid formation via aqueous two-phase systems**
Spheroids formed via aqueous two-phase systems (ATPS) were grown in a non-adhesive 96-well round bottom plate following previously published techniques using a robotic liquid handler (Gilson PipetMax, Mandel, Guelph ON)[1,2]. Briefly, a 0.2% (w/v) solution of Pluronics F108 in PBS was pipetted into each well and incubated for 1 hour at room temperature (23°C). The solution was aspirated, and the wells rinsed with reverse osmosis (RO) water before air drying. Plates were sterilized under UV light for 45 minutes prior to use. Stock solutions of 6% (w/v) polyethylene glycol (PEG) in complete media; and diluted to 5.4% in water prior to use. A cell-laden dextran (DEX) solution was prepared by mixing 85 µL of a 15% (w/v) dextran in PBS solution with 15 µL of a 17 x $10^6$ cells/mL suspension of HS-5 fibroblasts. To incorporate NiPAAM microgels into the spheroids, 1-3 µL of the functionalized microgel suspension was mixed into the cell-laden dextran depending on the desired microgel to spheroid ratio. 50 µL of the PEG solution was dispensed into each well of the non-adhesive 96 well-plate, and 1 µL of cell-laden DEX was carefully dispensed slightly above the bottom of each well. The plates were carefully transferred to a cell culture incubator (5% $CO_2$, 37 °C) for 1 hour before adding 75 µL of complete media and growing the spheroids for two days.

**Spheroid formation via micropocket hydrogel cavities**
Spheroids were formed in polyacrylamide micropockets using previously published protocols[3]. Polyacrylamide micropockets were cast using the 12 % acrylamide/0.24% bis-acrylamide formulation. Approximately 125 µL of the prepolymer polyacrylamide solution was dispensed over a 3D printed mold containing ~200 spherical structures across the surface area of a 12 mm coverslip to generously fill the mold. An MPS-treated 18 mm coverslip was placed on top of the mold, and the hydrogel was allowed to polymerize for 10 minutes. The polymerization grafted the polyacrylamide hydrogel to the coverslip, which was then gently separated from the 3D

printed mold, and washed 3 times in PBS. Gels were stored at 4 °C in PBS to equilibrate overnight, and sterilized under UV light for 45 minutes. PBS was aspirated prior to loading the micropocket gels with cells. A mixture containing 100 µL of a 15 x $10^6$ cells/mL suspension of the desired cell type with 1 µL of functionalized µTAM suspension was distributed over each hydrogel. The cells were left to settle into the micropockets for 5 minutes before submerging the entire polyacrylamide micropocket device in complete media. Spheroids then formed over 2 days in a standard cell culture incubator (5% CO2, 37 °C).

**Mouse breast cancer model**

Mice were housed at the Goodman Cancer Research Center animal facility where all procedures were performed in accordance with the animal care guidelines after obtaining ethics approval from the Animal Resource Centre of McGill University. For each replicate, a set of 4 female BALB/c mice (Charles River) at 8-10 weeks of age were randomly allocated a condition (sham, week 1, week 2 or week 3). Mice were anesthetized under isoflurane gas, as the 4th and the 9th mammary fat pads were injected using a 22G needle (Becton Dickinson) attached to a Hamilton syringe. Each gland was injected with a suspension of mCherry-labelled 4T1 cells at 5 x $10^5$ cells/mL with different concentrations of µTAMs in 25 µl of sterile PBS. The 4T1 tagged cells were generated using lentivirus and the lentivector pWPI-mCherry. Sham condition mice were injected only with a suspension of µTAMs in 25 µl of sterile PBS and left for 3 weeks

Mice were euthanized by cervical dislocation under isofluorane anesthesia. At the indicated time points, injected mammary fat pads or tumors were surgically isolated and immediately rinsed in sterile PBS.  Tumors exceeding 5 mm in thickness were sectioned to layers 4 ±1 mm in thickness to facilitate bead visualization and rinsed 10 times in sterile PBS. Tissue was immersed in sterile PBS in a 2-well chambered coverglass (Nunc™ Lab-Tek™) for immediate imaging.

**Temperature-controlled imaging and µTAMS size analysis**

Polyacrylamide phantoms, and multicellular spheroids were mounted in a Chamlide imaging chamber and submerged with 300 µL of PBS before being placed on a controlled stage warmer (Ibidi). Images were taken on an Olympus IX-73 microscope under epifluorescence (Olympus, X-CITE 120 LED), with an sCMOS Flash 4.0 Camera and Metamorph software (version 7.8.13.0), and automated stage (Zaber) to record and return to specified positions. Samples were mounted in a live-cell imaging chamber (Ibidi), and imaged initially at 37°C and during cool-down to room temperature at 30 minute intervals to ensure temperature equilibration and complete sensor size change. Live mouse tissue explants were imaged with an LSM700 laser scanning confocal microscope with a 20 x 0.8NA objective lens and ZEN software (Zeiss) in a temperature-controlled environmental chamber. The tissue was then incubated at 37˚C for 1 hour, and the same positions were re-imaged using the same imaging parameters. Images were deconvolved using the iterative deconvolve 3D plugin [4] and a point spread function generated by imaging 0.19 µm green TFM beads in identical imaging conditions as the µTAMs.

µTAMs that were damaged (missing chunk or fragment), clustered in a group, or partially exposed outside of given tissue were excluded from measurements. µTAMs that were less than 10 µm in diameter were excluded from stiffness analysis to reduce measurement error. All µTAM images were analyzed in FIJI by manually drawing a fitted ellipse around the µTAM and measuring the Feret's diameter for the microgel size, as well as shape descriptors for the circularity of the µTAM which was calculated within the software as:

$$circularity = \frac{4\pi \times Area}{\sqrt{Perimeter}}$$

**Shear rheometry for bulk mechanical characterization of hydrogels.**

The stiffness of each polyacrylamide and PNiPAAM gel formulation was measured using a parallel plate shear rheometer (Anton-Paar, MCR 302) in strain-controlled mode. Hydrogels fabricated for shear rheology were made by sandwiching 113 µL of the complete pre-gel solution (compositions provided in Supplementary Table S2) between two 12 mm coverslips treated with MPS as described earlier. After 10 minutes, the sandwiched polymerized hydrogels were placed in a multi-well plate and submerged in PBS. After three washes, the gels were left to swell overnight at 4°C. During testing, excess PBS was dried off the top and bottom of the samples, and adhesively mounted between rheometer plates. Storage and loss moduli were recorded over a strain sweep that was run from 1 to 50% at 10 Hz and verified to be plateau within this range. The moduli values were reported as an average of all the readings. Young's modulus (E) was calculated using $E = 2G(1 + \nu)$ where G is the average storage modulus, and ν is the Poisson's ratio of the hydrogel which was assumed to be 0.5 based on literature[5].

**Histology and staining**

Spheroids were fixed in 4% paraformaldehyde solution for at least 24 hours at 4°C. Spheroids in the micropockets were extracted and transferred using a clipped P1000 pipette tip placed directly over the chamber opening. Spheroids formed by ATPS were pipetted directly with a clipped pipette tip. Spheroids were collected into a 1.5 mL microcentrifuge tube with PBS embedded in paraffin blocks. Tissue blocks were sectioned at 4 µm and mounted on charged glass slides for histology.

For staining, the tissue sections were deparaffinised in xylene for 15 minutes and rehydrated using a decreasing ethanol gradient at 100%, 90% and 80% for 2-minute intervals. Slides were washed twice with PBS for 5 minutes, and permeabilized in 0.1% Triton-X solution for 5 minutes, before two additional PBS washes. Tissue sections were blocked with 1% BSA in PBS for 30 minutes at room temperature (23°C). An actin cytoskeletal and nuclear staining mixture of FITC-conjugated phalloidin (1 µg/mL) and hoescht 33258 (1 µg/mL) in 1% BSA was applied for 20 minutes. The slides were washed twice in PBS and once with water before coverslip mounting using Fluoromount Aqueous Mounting Media and sealing with clear nail polish.

**Histological Section Image Analysis**

All image analyses were performed using FIJI [6]. The cross-sectional area of the circular spheroids were segmented into 5 annuli of equal area. Cell density in each anulus was quantified with an automated nuclear count by thresholding the image to isolate the nuclei and performing a particle analysis count with a minimum of particle size of 20 µm$^2$. Nuclear orientation was analyzed by determining the difference between the expected angle for a circumferentially aligned nucleus ($\Theta_{expt}$) and the angle of the nucleus itself as determine by the particle analysis on FIJI. To get $\Theta_{expt}$, the angle at the center of a circle ($\Theta_r$) was calculated by taking the tangent angle between the X and Y distance of the nucleus to the center of the spheroid. $\Theta_{expt}$ was calculated by assuming spherical symmetry in the spheroid and taking the absolute value of $\Theta_r + 90°$ if $\Theta_r > 0°$ or $\Theta_r - 90°$ if $\Theta_r < 0°$.

**Statistical analysis**

Comparative data analysis of populations was performed without pre-specifying a required effect size. Datasets that were normally distributed, with similar variances between compared groups were analyzed using unpaired t-tests or two-way ANOVA to test for significance which was set at $\alpha= 0.05$. Post-hoc pairwise comparisons were conducted using the Bonferroni method. All statistical analyses were performed using GraphPad Prism v8.0.2 (San Diego, CA).

**Spatial modelling of measured stiffness distributions**

To construct a trend line that captures the envelope of stiffnesses observed into the depth of the multicellular spheroid structures, an inverse gaussian distribution function was selected and mathematically represented as:

$$f(x,\lambda,\mu,A,B,C) = A\bullet\left[\frac{\lambda}{2\pi(B+x)^2}\right]^{1/2}\bullet e^{-\frac{[\lambda(B+x)-\mu]^2}{2\mu^2(B+x)}} + C$$

Where ƒ represents the measured stiffness, and $x$ is the normalized distance towards the center of a spheroid. The main parameters in the inverse Gaussian distribution include the shape parameter of the distribution ($\lambda$) and the mean of the distribution (μ,). Parameters A, B, and C represent vertical scaling, horizontal shifting, and vertical shifting transformations to the original function. Best fit parameters were iterated to the highest $R^2$ value possible (Table S4) to fit the observed stiffness profile seen in our spheroids and describe the shape and location of stiffness peaks. Curve fitting and iterative optimization of model parameters to maximize the $R^2$ fit to data was performed in OriginLab 2018 Version 95E (Northampton, MA), using a non-linear fitting tool for the described user-defined functions.

**Finite element modelling of µTAM expansion**

Simulations were performed using the open-source software package FEBio[7] with the pre-strain plugin [8] to apply compressive loads to simulated µTAMs prior to release within an encapsulating matrix of defined stiffness. 3D spherical geometries were used to simulate the µTAMs (unit radius) embedded in a 10x larger encompassing sphere, to simulate an infinitely large matrix. The model was meshed with hexahedral elements, and a mesh size sensitivity analysis was performed. Less than 1% variation was observed in deformation for a mesh element size of 0.16 at the µTAM/matrix interface, for an r-ratio of 1.57. Fixed displacement boundary conditions were applied to the outer matrix surface, and a tied contact interface was defined at the µTAM/matrix interface. Linear elastic material properties and initial pre-strain of the µTAMs were defined and modulated based on experimental data. Analyses were conducted using a dynamic large deformation structural mechanical analysis, and data is reported as a fold change in µTAM size for matrices of various mechanical stiffness.
